# Novel structural insights on full-length human RAD52: Cryo-EM and beyond

**DOI:** 10.1101/2023.04.03.535362

**Authors:** Beatrice Balboni, Roberto Marotta, Francesco Rinaldi, Stefania Girotto, Andrea Cavalli

**Author notes:** These authors contributed equally (Beatrice Balboni, Roberto Marotta).

## Abstract

Human RAD52 is a DNA-binding protein involved in many DNA repair mechanisms and genomic stability maintenance. In the last few years, this protein was discovered to be a promising novel pharmacological target for anticancer synthetic lethality strategies since its inhibition or modulation, under specific genetic conditions, was proved to enhance therapies efficacy in various cancer cell types. Although the interest in RAD52 has exponentially grown in the last decade, most information about its structure and mechanism of action is still missing. This work provides novel insights into full-length RAD52 (RAD52 FL) protein, focusing on its structural and functional characterization. The Cryo-Electron Microscopy (Cryo-EM) structure of RAD52 FL, here presented at a resolution (2.16 Å) higher than the one currently available for RAD52 N-terminal X-ray structure, allows hypothesizing the role of individual amino acid residues. While the N-terminal region of RAD52 FL is structured in an undecameric ring, the C-terminal part is intrinsically disordered as fully characterized through SAXS and biophysical analyses. These detailed (atomic level) structural analyses will substantially impact future characterizations of RAD52 mechanisms of action and inhibitors development, particularly in the context of novel approaches to synthetic lethality.

## Introduction

RAD52 is a DNA-binding protein that is assumed to be involved in several DNA repair mechanisms such as Single Strand Annealing (SSA)^1–4^, Homologous Recombination (HR)^5–8^, stalled replication fork protection and processing^9,10^, RNA-dependent DNA repair^11–13^, which are activated under specific conditions throughout the cell cycle to control genome stability maintenance^11,14–16^. Nevertheless, human RAD52 has long been considered non-essential since it primarily plays an ancillary role in promoting RAD51-mediated DNA repair or serves as a backup mediator at specific cell cycle time points^17,18^. However, in recent years, many have been pieces of evidence demonstrating that RAD52 plays a primary role in specific cellular conditions, making this protein an attractive target for developing novel anticancer therapies. The decisive discovery occurred about ten years ago when it was observed that RAD52 depletion induces synthetic lethality in BRCA2, BRCA1, PALB2 mutated cancer cells^19–21^. Since then, many efforts have been made to structurally and functionally characterize human RAD52 to have a better understanding of its mechanism of action and to develop novel inhibitors for mono-or combination therapies.

RAD52 is a multimeric ring-shaped protein made of 418 amino acids. The N-terminal domain of the protein contains the oligomerization domain and a DNA binding domain^2,22,23^, whereas the C-terminal domain contains two binding sites for DNA repair protein RAD51 homolog 1 (RAD51) and a Replication Protein A (RPA), respectively^22,24^. For many years a heptameric ring conformation was ascribed to full-length RAD52 (RAD52 FL) based on indirect structural evidences^25–27^. Nevertheless, a recent cryo-EM structure (3.5 Å) from Kagawa and colleagues reported that RAD52 FL forms undecamers^28^. Notably, the structure of the N-terminal portion of the protein (residues 1-212) has been solved by X-ray (PDB ID: 1KN0). RAD52 N-terminal domain also organizes in a ring-shaped undecameric structure with a “mushroom-like” form, a “stem”, and a “domed cap”, and it has overall a deep positive charged cleft to accommodate DNA molecules wrapping around the protein ring (inner DNA binding site)^2,23,29^. Notably, a second DNA binding site (outer DNA binding site) is also reported ^29,30^.

This work presents the first cryo-EM structure of RAD52 FL (2.16 Å) at a resolution higher than that of the N-terminal domain via X-ray (2.85 Å). The present data allow, for the first time, a residue-by-residue analysis and comparison of RAD52 FL with other available structures. Through a multidisciplinary approach, cryo-EM, SAXS, computational simulation, and biophysical analyses, we confirmed, in agreement with Kagawa and colleagues, that full-length human RAD52 exists as an undecameric ring, matching the N-terminal domain organization, rather than the previously described heptameric arrangement ^28^. Furthermore, our analyses confirmed the extreme flexibility of the RAD52 C-terminal domain, which, as in its recombinant form, can be described as an intrinsically disordered protein. This observation further draws attention to the hitherto minor role played by the RAD52 C-terminal portion of the protein, which may assume a well-defined structure only upon partner binding.

## Materials and Methods

### Protein expression and purification

Histidine-tagged human RAD52 FL (RAD52 FL) expression vector (pET15b; *Addgene*) was transformed in *E. Coli* BL21 (DE3) Rosetta pLysS (*Sigma-Aldrich*). Freshly grown colonies were resuspended in Luria Broth (LB) buffer containing 100 ug/mL Ampicillin. Bacterial cultures were shaken at 180 rpm at 37°C to reach an optical density (OD_600_) of 0.8 and were then induced for 4 hours using 0.8 mM isopropil-β-D-1-tiogalattopiranoside (IPTG) at 25°C. Biomasses were finally harvested and put at -80°C.

Frozen pellets were thawed and resuspended in lysis buffer (25 mM TRIS-HCl pH 7.5, 500 mM NaCl, 5% glycerol, 10 mM Imidazole, Tween20 0.01%, 2mM DTT, Protease Inhibitors EDTA-free 1x (*Roche*)). The cell suspension was then lysed on ice by sonication and centrifuged for 1 hour at 30000g. The resulting surnatant was collected and loaded on a HisTrap HP chromatography column (*Cytiva*) equilibrated with Buffer A (25 mM TRIS-HCl pH 7.5, 500 mM NaCl, 5% glycerol, 10 mM Imidazole, Tween20 0.01%, 2mM DTT). Washing steps were performed with 4% and 8% of buffer B (25 mM TRIS-HCl pH 7.5, 500 mM NaCl, 5% glycerol, 0.5 M Imidazole, Tween20 0.01%, 2mM DTT). Protein fractions eluted at 40% buffer B and were pooled together and loaded on a 5 mL HiTrap Desalting chromatography column (*Cytiva*) equilibrated with buffer C (25 mM Tris-HCl pH 7.5, 150 mM NaCl, 5% glycerol, 1 mM DTT, 0.005% Tween20). The eluted protein was finally loaded on HiTrap Heparin HP chromatography column (*Cytiva*) equilibrated with buffer C. RAD52 FL pure protein eluted using a linear gradient up to 100% buffer D (25 mM Tris-HCl pH 7.5, 1.5 M NaCl, 5% glycerol, 1 mM DTT, 0.005% Tween20) in 5 CVs. The protein storage buffer was 25 mM Tris-HCl pH 7.5, 250 mM NaCl, 5% glycerol, 1 mM DTT, 0.005% Tween20. The purified protein was flask frozen in liquid nitrogen and stored at -80°C.

For histidine-tagged human RAD52 N-terminal domain (1-212), a similar expression and purification protocol was followed. Briefly, after the transformation and expression procedure as described above, the bacterial pellet was resuspended in a 50 mL volume of lysis buffer (25 mM TRIS-HCl pH 7.5, 500 mM NaCl, 5% glycerol, 10 mM Imidazole, Tween20 0.01%, 2mM DTT, Protease Inhibitors EDTA-free 1x (*Roche*). The cell suspension was then lysed on ice by sonication and centrifuged for 1 hour at 30000g. The collected surnatant was loaded on a 5 mL HisTrap HP chromatography column (*Cytiva*) equilibrated with Buffer A (25 mM TRIS-HCl pH 7.5, 500 mM NaCl, 5%glycerol, 10 mM Imidazole, Tween20 0.01%, 2mM DTT). Washing steps were performed with a 4%, 8%, and 40% of buffer B (25 mM TRIS-HCl pH 7.5, 500 mM NaCl, 5%glycerol, 0.5 M Imidazole, Tween20 0.01%, 2mM DTT). Fractions eluted at 100% buffer B (0.5 M Imidazole) were pooled in the following storage buffer: 25 mM Tris-HCl pH 7.5, 500 mM NaCl, 5% glycerol, 1 mM DTT, 0.005% Tween20. The purified protein was flask frozen in liquid nitrogen and stored at -80°C.

For oligomerization state analysis, RAD52 FL (20 μM) and RAD52 N-terminal (13.2 μM) (100 μL) were applied onto a size exclusion chromatographic column (Superdex200 Increase 3.2/300) previously equilibrated with buffer containing 20 mM Tris pH 7.4, 250 mM NaCl, 1% Glycerol. Runs were performed at 4°C through an ÄKTAmicro (GE Healthcare Life Sciences) system.

### Circular Dichroism

Far-UV CD spectra were recorded on a Jasco J-1100 spectropolarimeter (Jasco, Essex, United Kingdom), equipped with a temperature control system, using a 1 mm quartz cell. The spectra were recorded in the far-UV range 190–260 nm, using RAD52 FL and RAD52 N-terminal protein at 5 μM and 10 μM concentrations, respectively. Assay buffers were optimized to remove CD interference signals, avoiding the use of chlorine ions and glycerol and maintaining an ionic strength comparable with that of the storage buffers. RAD52 FL assay buffer was 25 mM phosphate buffer NaPi, pH 7.5 and 250 mM NaF; RAD52 N-terminal assay buffer was NaPi, pH 7.5 and 500 mM NaF. Constant N2 flush at 4.0 L/min was applied. Raw spectra were corrected for buffer contributions, and the detected signal was expressed as mean residue molar ellipticity [θ] (deg×cm2×dmol–1).

For protein secondary structure analysis, the scanning speed was set to 100 nm/min, digital integration time to 1 s, and the temperature set to 20 °C for all experiments. Each spectrum was obtained as an average of 10 scans. No shaking was applied to the samples during measurements. Data analysis was performed using *Dichroweb* (Institute of Structural and Molecular Biology Birkbeck College – University of London^31,32^).

For protein thermal stability analysis, CD experiments were performed using a temperature scan from 20°C to 95°C. No shaking was applied during data collection. Thermal stability was measured by monitoring the CD signal at 222 nm wavelength during the temperature scan.

### Dynamic light scattering (DLS)

RAD52 FL and RAD52 N-terminal protein samples were analyzed through DLS to test oligomerization propensity. Specifically, RAD52 FL and RAD52 N-terminal were tested, immediately after protein purification, in their storage buffers (25 mM Tris-HCl pH 7.5, 250 mM NaCl, 5% glycerol, 1 mM DTT, 0.005% Tween20, and 25 mM Tris-HCl pH 7.5, 500 mM NaCl, 5% glycerol, 1 mM DTT, 5 mM Imidazole, respectively) at 25°C, at 0.8 mg/mL (RAD52 FL) and 0.7 mg/mL (RAD52 N-terminal). Sizes of the samples were analyzed using Zetasizer Nanoparticles Analyzer Software (*Malvern*) using the standard operating procedures for size measurements, repeating the measurements scans 13 times for each sample. Collected data were expressed in terms of mass distribution. Reported data are the average of >3 independent experiments.

### SDS-Page

SDS-PAGE was performed using precast polyacrylamide gels (NuPAGE 4-12% BisTris Gel, *Invitrogen*). Different concentrations of protein samples were mixed with 4X Loading buffer (0.25 M Tris-HCl pH 6.8, 8% SDS, 0.3M DTT, 30% Glycerol, 0.4% Bromphenol Blue) before denaturation at 95°C for 5 minutes. After samples loading, precast gels were run in XCell SureLock Mini-Cell Electrophoresis System (*Invitrogen*) in MOPS SDS running buffer (*Invitrogen*) with a constant voltage of 120 mA for about 90 minutes. Gel was then stained with Comassie Blue staining buffer (40% EtOH, 10% Acetic Acid, 0.05% w/v comassie blue G-250) for 15 – 30 minutes and distained with a distaining Buffer (8% acetic acid, 25% EtOH). Protein bands images were visualized and quantified using ChemiDoc Imaging System (*BioRad*).

### Microscale Thermophoresis (MST)

MST measurements were performed using Monolith NT.115pico instrument (*NanoTemper Technologies*, Munich, Germany). Assays were conducted at 5%-10% (RED dye) LED excitation power and MST power of 40%. Premium capillaries from NanoTemper Technologies were used. Measurements were performed at 25 °C in the following buffer: 25 mM Hepes pH 7.5, 5% glycerol, 250 mM NaCl, 0.05% tween20.

The recombinant RAD52 FL protein was labeled with the Monolith labeling kit RED-NHS (ammine dye NT-647-NHS) according to manufacturer indications (*NanoTemper Technologies*). Before MST experiments, the labeled protein stocks were centrifuged at 20000 for 10 minutes to remove aggregates. The affinity parameter K_d_ was determined by performing the experiment in parallel on 16 capillaries, each containing a constant concentration of the labeled target (RAD52 FL, 10 nM) and increasing concentrations of unlabeled ligand (unlabeled RAD52 FL 0.7 nM 25 μM). The recorded MST data were then plotted as ΔF_norm_ against the ligand concentration to yield dose-response curves. Experiments were analyzed with MO.Control and MO.Affinity analysis software (NanoTemper Technologies).

### Fluorescensce Polarization (FP)

For fluorescence polarization experiments, 10 nM 6FAM (fluorescein)-conjugated dT_30_ ssDNA (*Merck*) was added to 16 different concentrations of RAD52 FL ranging between 6.25 μM and 0.2 nM in assay buffer (25 mM Tris-HCl pH 7.5, 62.5 mM NaCl, 5% glycerol, 1 mM DTT). 100 μL of each reaction was incubated in a flat black 96-plate (*Corning*) at room temperature for 15 minutes. FP of samples was then measured using a Spark Microplate multimodal reader instrument (*Tecan*) with the following setup Excitation wavelength 485 nm; Emission wavelength 535 nm; Emission Bandwidth 20 nm; Integration time 20 μs; reference mP value was set on 20 mP.

FP values were plotted as a function of protein concentrations and fitted using a sigmoidal dose-response curve with GraphPad Prism 7.0. Data are the average of multiple (>3) independent experiments.

### SEC-SAXS

SEC-SAXS experiments of RAD52 FL were performed at the BM29 beamline of the European Synchrotron Radiation Facility (ESRF, Grenoble, France)^33^. 100 µL of the sample were injected into a Superose 6 3.2/300 pre-equilibrated in 20 mM Tris pH 7.5, 250 mM NaCl, 1% glycerol, and the flow set to 0.075 mL/min. Data collection parameters are listed in Supplementary Table 1^34^. SEC-SAXS data were analyzed using Chromixs ^35^. Scattering frames corresponding to samples were selected, averaged, and subtracted from averaged buffer frames. The buffer-subtracted, 1D scattering curves were then processed using BioXTAS RAW to compute the radius of gyration, the dimensionless Kratky Plot, and obtain molecular mass estimates (Supplementary Table 1)^36^. Data were exported in .CSV format and re-graphed using GraphPad Prism 9 software. Considering the elevated number of N-terminal and C-terminal flexible residues to be modeled it was decided to generate small pools of 2000 models each through iterative cycles of RANCH (RANdom Chain generator) run via ATSAS online and utilizing the solved Cryo-EM structure as free-rigid body^37–39^. Input parameters are reported in Supplementary Table 1. Once 20000 models were created, these were locally pooled in a unique ensemble, and Form Factor MAKER (FFMAKER) was locally run^37–39^. The output file was then utilized for running Genetic Algorithm Judging the Optimization of the Ensemble (GAJOE)^37–39^. Model fittings, residuals, and Rg distributions were graphed using Graphpad Prism 9. SAXS Data have been deposited into SASBDB with accession number SASDQ49 – His-Tagged full-length DNA repair protein RAD52 homolog^40^.

### Negative Staining transmission electron microscopy

Negative staining experiments were performed on purified full-length RAD52 FL 0.1 mg/mL (buffer 25 mM TRIS-HCl pH 7.5, 250 mM NaCl, 5% glycerol, 0.005%Tween20). Briefly, a 5 μL drop of the sample was applied to previously plasma cleaned 400 mesh copper carbon film grids and stained with 1 wt/v % uranyl acetate solution. Data were collected on a JEM-1011 (JEOL) transmission electron microscope (TEM), with a thermionic source (W filament) and maximum acceleration voltage 100 kV equipped with Gatan Orius SC1000 CCD camera (4008 × 2672 active pixels) (see Supplementary Fig. 1A).

### Cryo-Electron Microscopy (Cryo-EM) CryoEM sample preparation and data collection

For cryo-EM grid preparation, a 3 µL droplet of purified RAD52 FL sample 0.7 mg/mL (in the following optimized buffer: 25 mM TRIS-HCl pH 7.5, 150 mM NaCl, 1% glycerol) was plunged frozen in liquid ethane cooled at liquid nitrogen temperature on glow discharged Quantifoil holey TEM grids (Cu, 300 mesh, 1.2/1.3 um) at 100 % humidity and 4.5° C. The grids were blotted with filter paper for 5 s using a Vitrobot Mark IV cryo-plunger (Thermo Fisher Scientific). Grid vitrification optimization was performed on a Tecnai F20 (Thermo Fisher Scientific) Schottky field emission gun transmission electron microscope equipped with an automated cryo-box, an Ultrascan 2kx2k CCD detector (Gatan), and a K3 direct electron detector (Gatan, Ametek).

Data screening and high-resolution data acquisition were performed at the EMBL Imaging Centre (Heidelberg, Germany) within the iNEXT project ID 15983. Data screening was performed on 831 gain-corrected counting mode movies acquired with a defocus ranging from -1.2 um to -2.5 um. Each movie, with a pixel size of 1.154 Å/pix, was composed of 34 frames collected with a total dose of 40.9 e^-^/Å^2^ (1.2 e^-^ /Å^2^/movie frame). This preliminary data set was acquired in EFTEM mode at 6.3 e^-^/pix/sec (8.6 sec/movie exposure) on a Thermo Fisher Scientific Glacios SelectrisX Falcon4 EC. The high-resolution data set consisted of 17,400 counting mode movies acquired with a defocus ranging from -0.8 um to 1.8 um. Each movie, with a pixel size of 0.731 Å/pix, was composed of 735 EER fractions collected with a total dose of 50 e^-^/Å^2^ (1 e^-^/Å^2^/ frame). The data set was acquired in EFTEM mode at 8 e^-^/pix/sec on a Thermo Fisher Scientific 300 kV cold FEG IC-Krios equipped with SelectrisX energy filter and Falcon4 EC direct electron detector. Data collection for both screening and high-resolution acquisitions has been performed using SerialEM software^41^ (see Supplementary Fig. 1B).

### Single particle image processing and 3D modeling

For the screening session motion correction was performed on all 831 movies (frames 2-31) with dose-weighting using the Relion3.1 MotionCor implementation^42^. CTF correction was performed with CTFFIND 4.1^43^ on all the 831-dose weighted motion-corrected micrographs. About 3900 representative particles were auto picked from 10 micrographs (391 particles/micrograph) using the Laplacian of Gaussian (LoG) filter. The obtained preliminary low pass filtered 2D class averages have then been used for automated particle picking. This resulted in 580952 particles extracted and down sampled for several iterative rounds of 2D classification and selection. A total of 258839 particles from 20 selected 2D classes were subjected to unsupervised 3D classifications (number of classes K = 4) using a RAD52 low-resolution initial model obtained from 2D averages with the 3D initial model algorithm implemented in Relion 3.1. Similar 3D classes have been obtained using a sphere as an unbiased initial model. The subsets of particles corresponding to the two more represented classes (class 3, 53% and class 4, 32%) corresponding to 221976 particles, after being re-extracted at full resolution, were used for the final refinement. The final RAD52 FL electron density map was resolved at 3.4 Å by the 0.143 FSC criterion after post-processing, CTF refinement, and polishing (Supplementary Table 2).

Motion correction and contrast transfer function (CTF) estimation were performed ‘on the fly’ during data acquisition for the high-resolution data collection. A preliminary particle manual picking was accomplished on about 30 selected micrographs (around 130 particles/micrograph) followed by a 2D classification on around 3600 extracted particles was performed to assess the overall sample quality and set up the correct particle box size. Both LoG filter parameters and box size optimizations were set up on 100 selected micrographs and then applied for particle auto-picking on all 14700 rescaled (1:2) micrographs. This resulted in 4124609 particles (281 particles/micrograph on average) extracted and down-sampled for several iterative round of 2D classification and selection. 2,325,722 particles from 65 selected 2D class averages were subjected to unsupervised 3D classifications (number of classes K = 4) using a RAD52 FL low-resolution initial model obtained from selected 2D class averages with the 3D initial model algorithm as implemented in Relion 4.0^44^. The subsets of particles corresponding to the more represented 3D classes (class 3 and 4 corresponding to 34% and 36% of particles, respectively), were used separately for the final refinements (Supplementary Fig. 2A). After CTF refinement and particle polishing, the best-resolved cryo-electron density map, resolved at 2.16 Å by the 0.143 FSC criterion, was obtained from 3D class 4 extracted particles, imposing c11 symmetry (Supplementary Fig. 2C, Supplementary Table 2). Compared to RAD52 FL map with c11 symmetry, the map obtained without imposing symmetry was similar although less resolved (Supplementary Fig. 2D, Supplementary Fig.s. 3A, C). The cryo-electron density maps obtained from 3D class 3, corresponding to more than 787000 particles, after CTF refinement and without imposing symmetry, was resolved at 2.6 Å by the 0.143 FSC criterion (Supplementary Fig.s. 2B, 3B, 3D). The human RAD52 FL protein model was obtained from the RAD52_25-208_ crystal structure (PDB ID 1KN0) after several iterative cycles of Phenix real space refinement^45^ and COOT^46^ manual adjustments. Cross-correlation analyses, measures of distances, model superimposition, 3D visualizations, and rendering were performed using Chimera^47^ and ChimeraX^48^.

## Results

### RAD52 full length has an undecameric ring-shape structure that forms high molecular weight superstructures

Human RAD52 FL (47 kDa) and its truncated N-terminal (1-212) domain were expressed and purified as recombinant proteins with a 6xHis tag at the N-terminal (Fig. 1A). Circular Dichroism (CD) analyses showed that RAD52 FL is characterized by similar thermal stability (Supplementary Fig. 4A) and a more unstructured profile than its N-terminal counterpart with 9% of alpha helices, 7% beta sheets, and 51% unordered regions (Fig. 1B).

**Figure 1.**
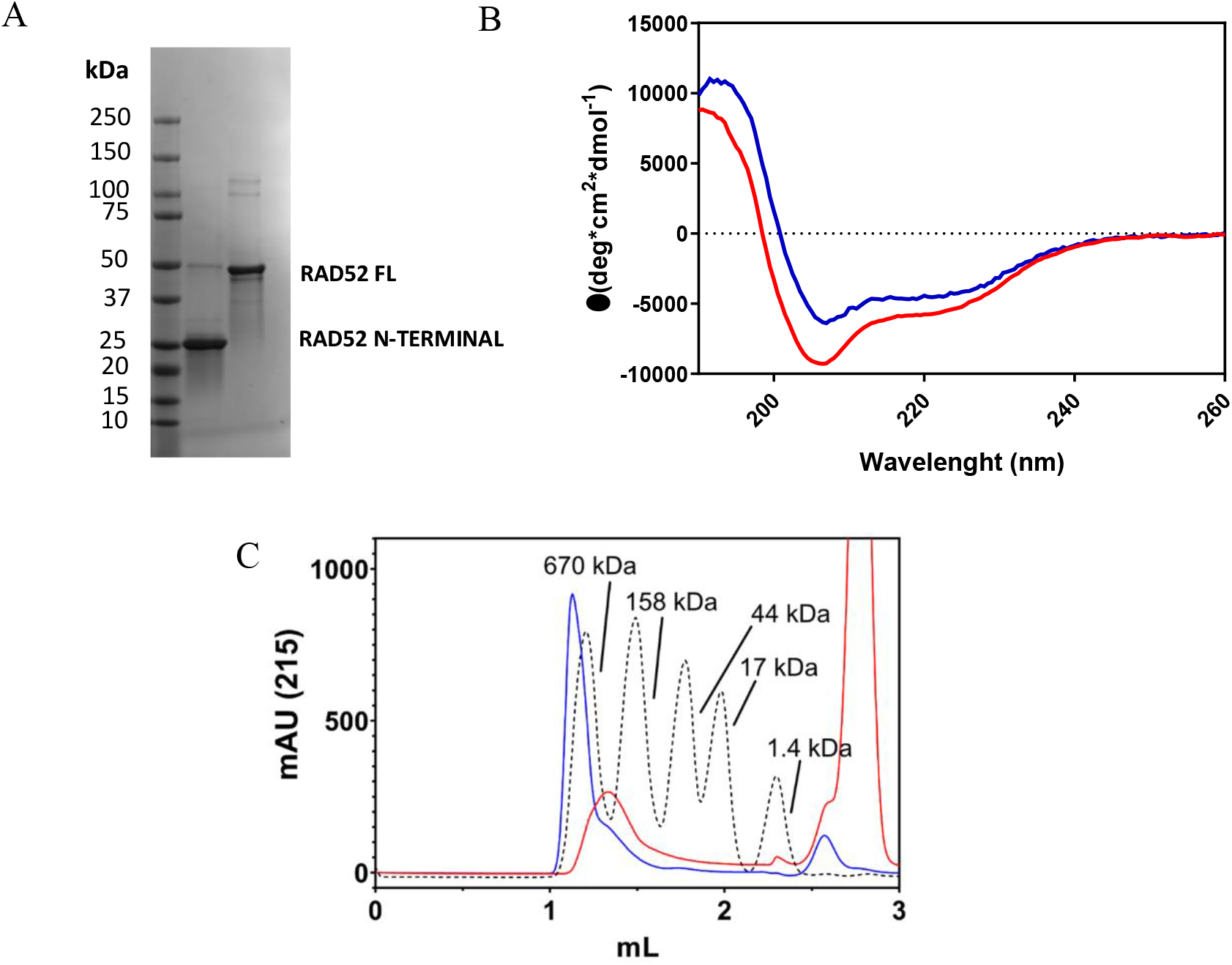
Characterization of recombinant RAD52 FL and N-terminal. 1) A: SDS-Page analysis of RAD52 N-terminal domain and FL samples; B: Secondary structure determination of RAD52 FL (red) and RAD52 N-terminal domain (blue) through CD spectra comparison; data analyses were performed after normalization; Dichroweb software (Birkberg college; University of London) was used to predict the secondary structure information; C: Comparison of RAD52 FL (red) and RAD52 N-terminal domain (1-212) (blue) SEC elution profiles; dashed lines represent the elution profiles of reference proteins.

Size exclusion chromatographic analysis showed that RAD52 FL forms high molecular weight (mw) species (Fig. 1C), whereas the N-terminal counterpart, although still able to form them, shows a lower propensity to do it. In addition, light scattering (DLS) analyses highlighted the formation of a highly polydisperse and heterogenic RAD52 FL sample with the most abundant species having an average hydrodynamic radius of 10.52 nm and an estimated average molecular weight of 828.9 kDa (Supplementary Fig. 4B), when tested at 0.8 mg/mL. DLS on the N-terminal domain still showed the formation of a polydisperse and heterogenic sample, but the most abundant species had an average smaller hydrodynamic radius (6.77 nm) and a lower estimated average molecular weight (295.5 kDa) roughly corresponding to an undecamer ring unit (Supplementary Fig. 4B). To further characterize the propensity to self-oligomerize of the FL protein microscale thermophoresis (MST) binding experiments were performed providing an apparent affinity constant of 13 μM (K_d_) (Fig. 2A). The highly complicated, partially unstructured and dynamic arrangement of RAD52 FL has so far prevented obtaining a 3D structure by X-ray crystallography. Nevertheless, a Cryo-EM approach was pursued, which is more suitable for gaining further structural insights into RAD52 FL sample.

**Figure 2.**
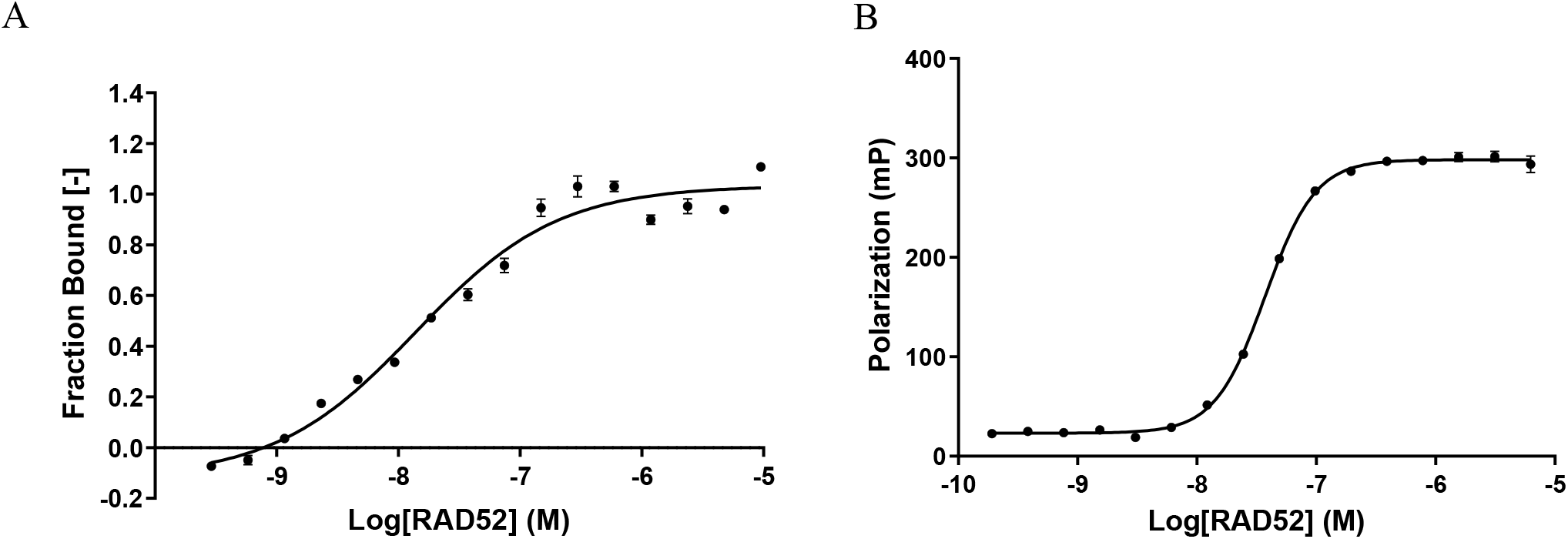
RAD52 FL self-oligomerization and DNA binding. A: MST analysis of protein-protein titration showing RAD52 FL tendency to self-oligomerize. The fitting curve was determined using non-linear regression (apparent K_d_ = 14.2 ± 4.0 nM). The data are the average of 2 replicates; B: Direct FP assay of ssDNA-RAD52 binding. The plot reports polarization data (mP) recorded upon addition of increasing RAD52 concentrations (Log[RAD52]) into a solution containing 10 nM 5’-FAM labeled ssDNA. The fitting curve was determined using non-linear regression (apparent K_d_ = 37.2 ± 0.4 nM).

RAD52 FL protein structure was obtained by Cryo-EM single particle analysis (Fig. 3A). The map was determined at 2.16 Å resolution imposing C11 symmetry, as we observed top views 2D class averages with a clear C11 symmetry (Fig. 3B and Supplementary Fig.. 5A, B). Our results pointed out that RAD52 FL protein complex, independently from the imposed symmetry, organizes in undecameric ring units, consisting in a mushroom-like closed ring composed of a steam region, formed by the β-β-β-α fold of each monomer (residues 79-156), and by a domed cup region that ends with a flat top (Fig. 3A)^23,29^. The stem region is quite rigid, except for some portions in loops L6 and L7. Instead of the β-hairpin loop (part of β-sheet β1, the loop L3, and part of β-sheet β2), a large portion of the L10 loop and small regions at the top of the domed cup, corresponding to the N- and C-terminal portions of RAD52 FL, were quite flexible, as pointed out by ResMap analysis (Supplementary Fig.. 5C). In our data collection, based on more than 2.3 million cleaned particles (see Materials and Methods), RAD52 FL protein complex was always organized in undecameric rings: no minor 2D or 3D classes corresponding to other ring organizations, have been observed (Fig.3A, B, Supplementary Fig.s. 2A, and 5A).

**Figure 3.**
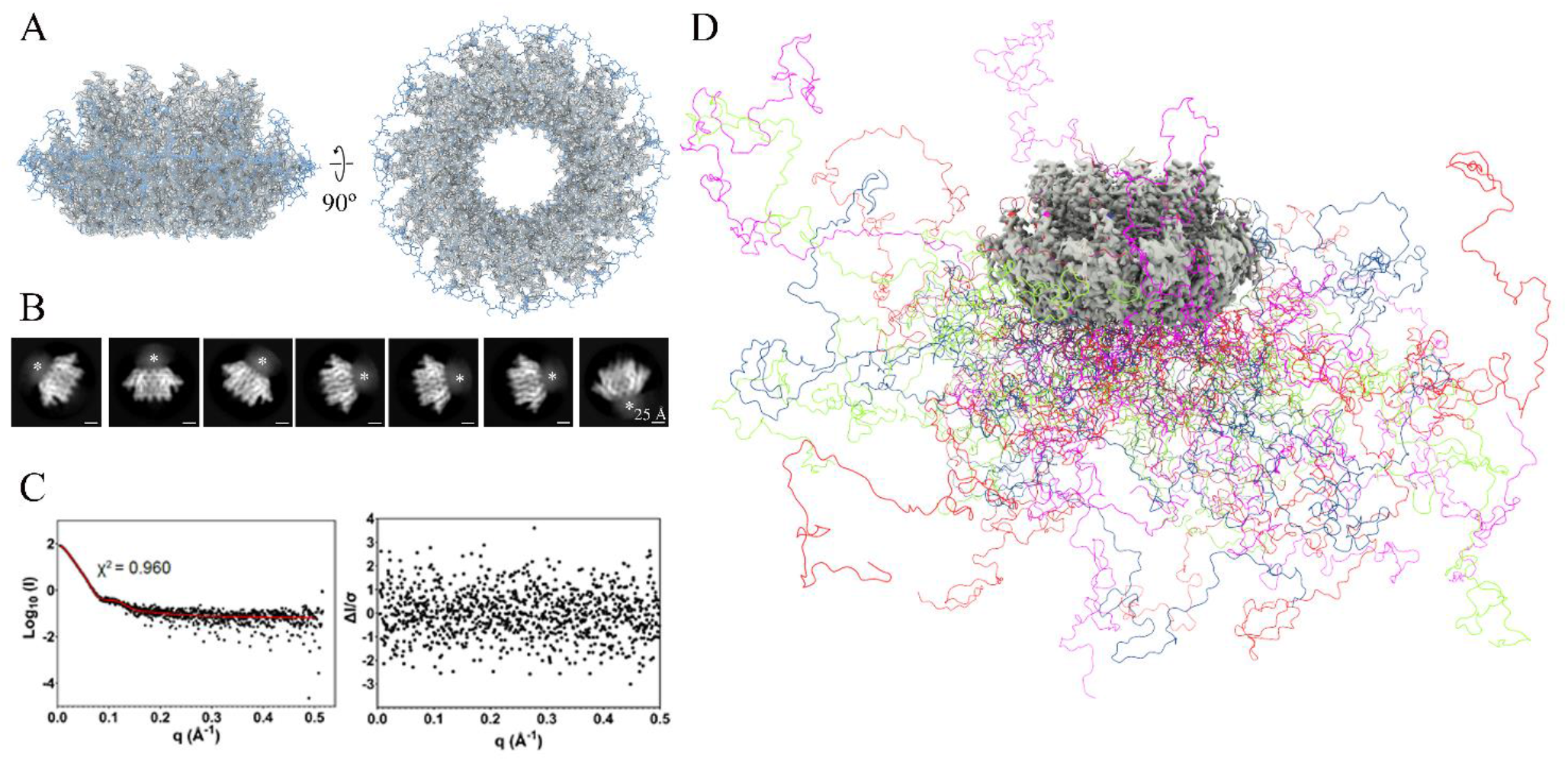
The C-terminal domain of RAD52 FL is an intrinsic disordered region (IDR). A: RAD52 FL cryo-EM density map resolved at 2.16 Å fitted with the model derived from the crystal structure of the RAD52 N-terminal truncated crystallographic model (PDB ID 1KN0^28^) in side (left) and top (right) views. B: representative unsupervised 2D class averages in side views showing an undefined electron density cloud close to the top of the ring (asterisks). C: Left: model fit to experimental SAXS data. Right: model residuals. D: in different colors the overlay of the four most representative SAXS models obtained by modelling the N-terminal and C-terminal domains with EOM and utilizing the Cryo-EM structure as rigid body.

A similar undecameric ring arrangement has been observed for RAD52 N-terminal domain^2,23^. Fluorescence polarization (FP) experiments show an apparent affinity constant of 37.2 nM (K_d_) for DNA binding to RAD52 FL, which is very close to the one already reported in the literature (∼6 nM) for RAD52 N-terminal (Fig. 2B), thus supporting a significant similarity between the two proteins^29^.

In the present cryo-electron density map, most amino acid side chains between Ser25 and Cys208 are well resolved and clearly recognizable (Fig. 3A). The map at both the N-(residues 1-22) and C-terminus (residues 209-430) are instead largely missing (Fig. 3A). Compared to the available RAD52 structures, in our model Val23 and Leu24 at the N-terminal and Arg209 (only in chains B, F, I and K) at the C-terminal are well observable (Supplementary Fig.. 6). Undefined electron dense clouds, close to the top of the rings, were visible in the majority of the 2D class averages side views (Fig 3B, and Supplementary Fig.. 5A); large unstructured regions were present, in a correspondent position and at lower sigma values, in the two best resolved 3D class averages (Supplementary Fig.. 7A, B).

### C-terminal domain of RAD52 FL is an intrinsically disordered region (IDR)

AlphaFold^49^, flDPnn^50^, PONDR-FIT^51^, and IUPred^52,53^ predicted that a large portion of the RAD52 FL C-terminal domain (residues 211 to 418) is unstructured and disordered (Supplementary Fig.. 8) in agreement with the cryo-EM data above reported. Size Exclusion Chromatography coupled with Small-Angle Synchrotron X-ray Scattering (SEC-SAXS) (SASBDB ID SASDQ49) experiments were then pursued to further confirm the disorder of RAD52 FL C-terminal domain and to characterize its behavior in solution. The primary data analysis showed a molecular weight of roughly 600 kDa, in agreement with previously reported DLS data, considering the concentration-dependent formation of high mw superstructures and sample dilution occurring during SEC (performed before SAXS data collection). SAXS data analysis also allowed us to calculate the radius of gyration (Rg) of 81.28 Å (8.128 nm) (Supplementary Fig.. 9). Dimensionless Kratky plot confirmed that the protein was folded but partially behaving as a flexible system supporting previous findings obtained by comparing RAD52 N-terminal domain with RAD52 FL by CD and Cryo-EM. Specifically, in this representation, a shift of the peak-maximum compared to the Guiner-Kratky point (red crosshairs) and a plateau at high q values not decreasing to 0 was reported. These features are typical of proteins harboring flexible portions^54–56^ (Fig. 3C and Supplementary Fig.. 9). Also, the p(r) function, calculated through the Indirect Fourier Transformation (IFT), showed a smooth decrement to 0 and large Dmax of the analyzed particles indicative of protein harboring flexible domains (Supplementary Fig.s. 9, 10)^55,57^. Based on the results obtained from Cryo-EM and computational approaches, the N-terminal (amino-acids 1-36, comprising the His-Tag) and the C-terminal domains (amino-acids 223-432, comprising the His-Tag)^57,58^ were modeled as disordered regions (Fig. 3D, Supplementary Fig.. 10). The modeling excellently fitted the experimental data with a χ2 of 0.960, and the overlay of the four most representative models allow to appreciate the extreme flexibility of the C-terminal domain in solution (Fig. 3D).

### Arg55 may act as a “gate” for DNA binding in RAD52 inner site

The presented RAD52 FL model (PDB ID 8BJM) is very similar to the other RAD52 published structures, with average root-mean-square deviations (RMSD) ranging from 0.539 to 0.671 (see Supplementary Table 3). In all RAD52 models, the largest Cα to Cα distances were observed mainly in the β-hairpin region loop L3, and to a lesser extent, in the loops L10, L8, and L6 (Fig. 4 A-C). Concerning loops L3 and L10, the largest average Cα to Cα distances were observed between our model and the N-terminal truncated crystallographic model (PDB ID 1KN0^29^). The largest Cα to Cα distances in loops L6 and L8 were instead observed between our model and the RAD52 outer (PDB ID 5XS0^29^) and inner (PDB ID 5XRZ^29^) DNA binding site models, respectively (Fig. 4 A-C).

**Figure 4.**
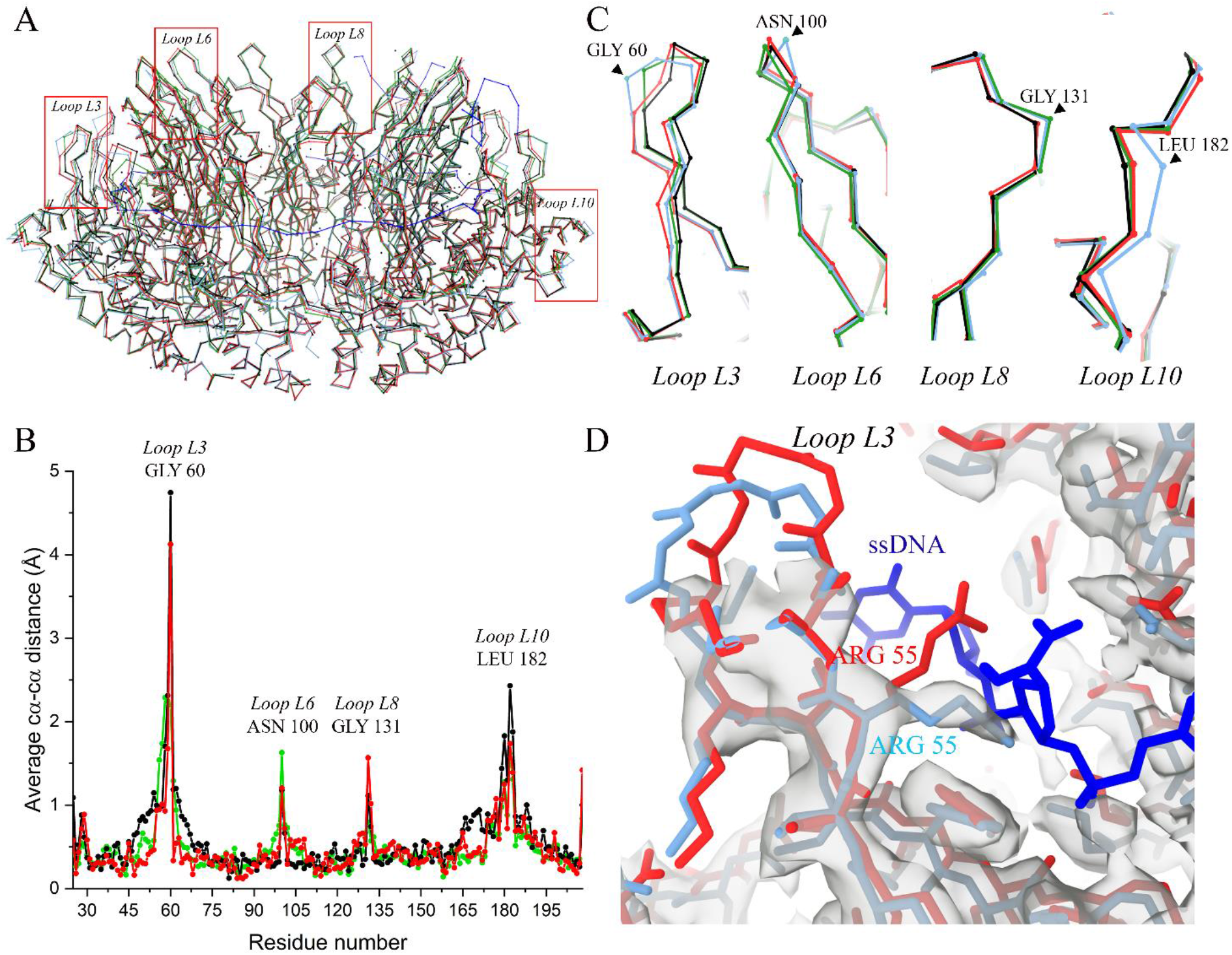
Comparison of RAD52 FL with the crystal structure of RAD52 N-terminal domain. A: superimposed average Cα traces of the RAD52 FL model with the RAD52_25-208_ model alone (PDB ID 1KN0^22^) and the inner (PDB ID 5XRZ^28^) and outer (PDB ID 5XS0^28^) DNA binding site models. B: graph showing the average Cα-Cα distances among the models shown in A. C: expansions of the boxed regions in A; D: detail of the RAD52 FL β-hairpin loop portion containing Arg55 (Loop L3) fitted in the cryo-EM electron density map superimposed to same region from the crystal structure of the RAD52_25-208_ inner DNA binding site model (PDB ID: 5XRZ^28^). Color code: the RAD52 FL, the RAD52_25-208_ model alone (PDB ID 1KN0^22^), the inner (PDB ID:5XRZ^28^) and outer (PDB ID 5XS0^28^) site DNA binding models are in cyan, black, red and green, respectively. The ssDNA is in blue.

Among the residues essential for ssDNA binding at the inner binding site (PDB ID 5XRZ^29^), in our structure, only Arg55 (hairpin loop, the last residue of β-sheet β1), significantly differs in position (Fig. 4D). Indeed, after model alignment, in all chains of our model Arg55 potentially clashes with the phosphate-deoxyribose backbone of the ssDNA (Fig. 4D). The other residues involved in ssDNA inner site interaction, i.e., Lys152, Arg153, and Arg156, all belonging to helix α3, did not significantly differ in position (not shown). Our cryo-electron density map was instead less defined at the residues involved in the ssDNA interaction in the outer DNA binding site, i.e. Lys102 (β-sheet β4) and Lys133 (Loop L8). Lys102, in the majority of chains, was apparently farther from the ssDNA phosphate groups than the correspondent residue in the outer DNA binding site model (PDB ID 5XS0^29^) (Supplementary Fig.. 11). These data suggest that in the reported cryo-EM structure Arg55 is not available for DNA binding at the inner DNA binding site.

## Discussion

This work presents the first high-resolution cryo-EM structure (2.16 Å) of the full-length human RAD52, together with its SEC-SAXS and biophysical characterizations. Our extensive single particle analysis unambiguously shows that RAD52 FL protein forms ring-shaped undecameric structures, closely matching the conformation of the crystallographic RAD52 N-terminal domain (PDB ID 1KN0^23^). Indeed, the observed largest Cα to Cα distances among our model and the crystal structures of the RAD52 N-terminal domain match substantially the flexible regions in the protein complex (Fig. 4). These data are also supported by similar DNA binding affinity constant for RAD52 FL and RAD52 N-terminal as recorded by FP. These results contrast with previous low-resolution data that inferred a heptameric ring arrangement rather than an undecameric one for the FL form^7,23,26,59^. Nevertheless, extensive data analysis excludes the coexistence of minor complexes with different symmetries in our samples.

A detailed residues analysis of the N-terminal domain proved the importance of Arg55 as a key residue for RAD52 DNA binding activity: Arg55 is known to have an essential role for RAD52 ssDNA binding since it anchors the ssDNA to the RAD52 inner DNA binding site^29^. Once the ssDNA is bound to the inner DNA binding site, Arg55 has been proposed to uniquely act as an entry/exit gate for ssDNA during homology search^29^. The position of Arg55 in our model is clearly not compatible with the binding of the ssDNA to the inner DNA binding site. Its position should change to allow the ssDNA to bind to this site. Thus, Arg55 may also work as an entry/exit gate for ssDNA in the first phases of RAD52-ssDNA interaction, not only during homology search. This result is also coherent with a milder effect on the DNA annealing activity observed for RAD52 FL R55A mutation^29^: alanine in position 55 allows the correct positioning of the ssDNA to the inner DNA binding site, but then it is too small for the ssDNA partial ejection, essential for homology search.

The residues involved in the ssDNA interaction at the outer DNA binding site are pretty flexible, as their corresponding electron density map is poorly defined. In our model, Lys102 was too far from the ssDNA phosphate groups to allow the formation of hydrogen bonds and stabilize the ssDNA^29^. Conformational changes in this region, which is indeed flexible, are probably needed to allow ssDNA binding.

In the final cryo-EM map, the nearly complete absence of the RAD52 C-terminal domains, putatively visible as an undefined electron density cloud in both 2D and 3D single particle analysis classifications, suggests that the RAD52 C-terminal regions occupy multiple positions and are highly disordered and flexible^60^. These results agree with CD data highlighting that RAD52 FL displays a more disordered profile than the RAD52 N-terminal domain alone and are further corroborated by computational analyses^61,62^. Additionally, SEC-SAXS analyses of RAD52 FL strengthen this structural evidence providing novel insights into the extreme flexibility of the RAD52 C terminal domain.

In light of these results, we propose that the human RAD52 FL is an intrinsically disordered protein (IDP) that displays two regions with completely different functions: a largely structured N-terminal domain and an intrinsically disordered region (IDR) of similar size (i.e., about 200 residues), corresponding to its C-terminal domain^63^. While the N-terminal region of RAD52 exerts its DNA binding function, the C-terminal domain could be required to find other RAD52 ring-shaped structures and help the DNA strand homology search and annealing^64,65^.

Indeed, RAD52 FL, with respect to its N-truncated counterpart, has a higher tendency to form high mw superstructures, meaning that the C-terminal part of the protein has an effect and likely a physiological function in the formation of larger complexes. Moreover, the RAD52 C-terminal IDR mediates the interaction with RPA and RAD51-ssDNA complex ^1,22,24,25^, having specific binding sites for both proteins^24,66^. RAD52 is known to associate with RAD51-DNA complexes forming globular structures along RAD51 nucleoprotein filament^24,25^. Noteworthy, the IDRs are often involved specifically in molecular recognition interactions, being able to wraps-up and bind several partners simultaneously by structural accommodation at the binding interfaces, with faster rates of association and dissociation^67^. We speculate that the RAD52 FL C-terminal region could bind other RAD52 FL rings, link both RPA-ssDNA and RAD51-ssDNA, and quickly detach from its partners during presynaptic complex assembly^68^.

The complex and dynamic RAD52 arrangement in high molecular weight superstructures frequently hampers its biophysical characterization. In fact, in this manuscript, only apparent parameters were reported due to simultaneous binding and oligomerization events. Nevertheless, in the dynamic nature of this protein lays its real physiological role and value. A thorough understanding of RAD52 self-association and DNA binding mechanism that is supported, and most likely fostered, by its oligomerization will be critical to unveil RAD52 mechanism of action.

Although the characterization of the structure and mechanism of action of RAD52 is not completely clarified and beyond the scope of this paper, we believe that this work can lay the foundation for further studies and for a better understanding of the physiological and pathological behavior of RAD52 FL in the presence of protein interactors and DNA. These analyses pave the way for the structural characterization of RAD52 in the presence of either protein interactors or DNA and its physiological mechanism of action, which may be related to high mw superstructures formations. In the coming years, understanding the protein behavior in the presence of intracellular partners will be crucial for developing novel RAD52 inhibitors, which could burst novel synthetic lethality-based anticancer therapies.

## Acknowledgments

This work has been supported by iNEXT-Discovery, project number 22228, funded by the Horizon 2020 program of the European Commission. We acknowledge the access and services provided by the Imaging Centre at the European Molecular Biology Laboratory (EMBL IC), generously supported by the Boehringer Ingelheim Foundation. We thank Dr. Simone Mattei (EMBL Imaging Centre) and Dr. Simon Fromm (EMBL Imaging center) for their support in data collection. The authors thank the BioSAXS-BM29 at European Synchrotron Radiation Facility (ESRF) for providing the synchrotron radiation facility for SAXS measurements. The authors thank Luigi Scietti and Giuseppe Ciossani of the IEO Biochemistry and Structural Biology Unit for the fruitful discussions.

## Fundings

We thank the Italian Institute of Technology for supporting and financing research activities in the field of synthetic lethality and anticancer drug discovery. This work was further supported by the Italian Association for Cancer Research (AIRC) through Grant IG 2018 Id.21386.

## Notes

### Competing Interest Statement

The authors have declared no competing interest.

## References

1. Grimme, J. M. et al. Human Rad52 binds and wraps single-stranded DNA and mediates annealing via two hRad52-ssDNA complexes. Nucleic Acids Res 38, 2917–2930 (2010).

2. Singleton, M. R., Wentzell, L. M., Liu, Y., West, S. C. & Wigley, D. B. Structure of the single-strand annealing domain of human RAD52 protein. Proc Natl Acad Sci U S A 99, 13492–13497 (2002).

3. Reddy, G., Golub, E. I. & Radding, C. M. Human Rad52 protein promotes single-strand DNA annealing followed by branch migration. Mutation Research - Fundamental and Molecular Mechanisms of Mutagenesis 377, 53–59 (1997).

4. Bhargava, R., Onyango, D. O. & Stark, J. M. Regulation of Single Strand Annealing and its role in genome maintenance Chromosomal break repair by the Single Strand Annealing (SSA) pathway. Trends Genet 32, 566–575 (2016).

5. Sugiyama, T., Kantake, N., Wu, Y. & Kowalczykowski, S. C. Rad52-mediated DNA annealing after Rad51-mediated DNA strand exchange promotes second ssDNA capture. EMBO J 25, 5539–5548 (2006).

6. New, J. H., Sugiyama, T., Zaitseva, E. & Kowalczykowski, S. C. Rad52 protein stimulates DNA strand exchange by Rad51 and replication protein A. Nature 391, 407–410 (1998).

7. Hanamshet, K., Mazina, O. M. & Mazin, A. v. Reappearance from obscurity: Mammalian Rad52 in homologous recombination. Genes (Basel) 7, 1–18 (2016).

8. Mahajan, S., Raina, K., Verma, S. & Rao, B. J. Human RAD52 protein regulates homologous recombination and checkpoint function in BRCA2 deficient cells. International Journal of Biochemistry and Cell Biology 107, 128–139 (2019).

9. Zeman, M. K. & Cimprich, K. A. Causes and consequences of replication stress. Nat Cell Biol 16, 2–9 (2014).

10. Kondratick, C. M., Washington, M. T. & Spies, M. Making Choices: DNA Replication Fork Recovery Mechanisms. Semin Cell Dev Biol 113, 27–37 (2021).

11. Rossi, M. J., Didomenico, S. F., Patel, M. & Mazin, A. v. RAD52 : Paradigm of Synthetic Lethality and New Developments. 12, 1–16 (2021).

12. McDevitt, S., Rusanov, T., Kent, T., Chandramouly, G. & Pomerantz, R. T. How RNA transcripts coordinate DNA recombination and repair. Nat Commun 9, 1–10 (2018).

13. Mazina, O. M., Keskin, H., Hanamshet, K., Storici, F. & Mazin, A. v. Rad52 Inverse Strand Exchange Drives RNA-Templated DNA Double-Strand Break Repair. Mol Cell 67, 19–29.e3 (2017).

14. Jalan, M., Olsen, K. S. & Powell, S. N. Emerging roles of RAD52 in genome maintenance. Cancers (Basel) 11, (2019).

15. Toma, M., Sullivan-Reed, K., Śliwiński, T. & Skorski, T. RAD52 as a potential target for synthetic lethality-based anticancer therapies. Cancers (Basel) 11, (2019).

16. Nogueira, A., Fernandes, M., Catarino, R. & Medeiros, R. RAD52 functions in homologous recombination and its importance on genomic integrity maintenance and cancer therapy. Cancers (Basel) 11, (2019).

17. Yamaguchi-Iwai, Y. et al. Homologous recombination, but not DNA repair, is reduced in vertebrate cells deficient in RAD52. Mol Cell Biol 18, 6430–6435 (1998).

18. Rijkers, T. et al. Targeted inactivation of mouse RAD52 reduces homologous recombination but not resistance to ionizing radiation. Mol Cell Biol 18, 6423–6429 (1998).

19. Lok, B. H., Carley, A. C., Tchang, B. & Powell, S. N. RAD52 inactivation is synthetically lethal with deficiencies in BRCA1 and PALB2 in addition to BRCA2 through RAD51-mediated homologous recombination. Oncogene (2013) doi:10.1038/onc.2012.391.

20. Feng, Z. et al. Rad52 inactivation is synthetically lethal with BRCA2 deficiency. Proc Natl Acad Sci U S A 108, 686–691 (2011).

21. Sullivan-Reed, K. et al. Simultaneous Targeting of PARP1 and RAD52 Triggers Dual Synthetic Lethality in BRCA-Deficient Tumor Cells. Cell Rep (2018) doi:10.1016/j.celrep.2018.05.034.

22. Kagawa, W., Kurumizaka, H., Ikawa, S., Yokoyama, S. & Shibata, T. Homologous Pairing Promoted by the Human, Rad52 Protein. Journal of Biological Chemistry 276, 35201–35208 (2001).

23. Kagawa, W. et al. Crystal structure of the homologous-pairing domain from the human Rad52 recombinase in the undecameric form. Mol Cell 10, 359–371 (2002).

24. Ranatunga, W. et al. Human RAD52 Exhibits Two Modes of Self-association. Journal of Biological Chemistry 276, 15876–15880 (2001).

25. van Dyck, E., Stasiak, A. Z., Stasiak, A. & West, S. C. Visualization of recombination intermediates produced by RAD52-mediated single-strand annealing. EMBO Rep 2, 905–909 (2001).

26. Stasiak, A. Z. et al. The human Rad52 protein exists as a heptameric ring. Current Biology 10, 337–340 (2000).

27. van Dyck, E., Hajibagheri, N. M. A., Stasiak, A. & West, S. C. Visualisation of human Rad52 protein and its complexes with hRad51 and DNA. J Mol Biol 284, 1027–1038 (1998).

28. Kinoshita, C. et al. The cryo-EM structure of full-length RAD52 protein contains an undecameric ring. FEBS Open Bio (2023) doi:10.1002/2211-5463.13565.

29. Saotome, M. et al. Structural Basis of Homology-Directed DNA Repair Mediated by RAD52. iScience 3, 50–62 (2018).

30. Lobley, A., Whitmore, L. & Wallace, B. A. DICHROWEB: an interactive website for the analysis of protein secondary structure from circular dichroism spectra. Bioinformatics 18, 211–212 (2002).

31. Miles, A. J., Ramalli, S. G. & Wallace, B. A. DichroWeb, a website for calculating protein secondary structure from circular dichroism spectroscopic data. Protein Science 31, 37–46 (2022).

32. Tully, M. D. et al. BioSAXS at European Synchrotron Radiation Facility –Extremely Brilliant Source: BM29 with an upgraded source, detector, robot, sample environment, data collection and analysis software. J Synchrotron Radiat 30, 258–266 (2023).

33. Rinaldi, F., Hočevar, J. & Scietti, L. Proteins related to pathogenesis of diseases, viral proteins, cell division, signalling and chromatin processes [Dataset]. European Synchrotron Radiation Facility (2025) doi:https://doi.org/10.15151/ESRF-ES-771426690.

34. Panjkovich, A. & Svergun, D. I. CHROMIXS: automatic and interactive analysis of chromatography-coupled small-angle X-ray scattering data. Bioinformatics 34, 1944–1946 (2018).

35. Hopkins, J. B., Gillilan, R. E. & Skou, S. BioXTAS RAW: Improvements to a free open-source program for small-angle X-ray scattering data reduction and analysis. J Appl Crystallogr 50, 1545–1553 (2017).

36. Tria, G., Mertens, H. D. T., Kachala, M. & Svergun, D. I. Advanced ensemble modelling of flexible macromolecules using X-ray solution scattering. IUCrJ 2, 207–217 (2015).

37. Bernadó, P., Mylonas, E., Petoukhov, M. V., Blackledge, M. & Svergun, D. I. Structural characterization of flexible proteins using small-angle X-ray scattering. J Am Chem Soc 129, 5656–5664 (2007).

38. Bernadó, P. Effect of interdomain dynamics on the structure determination of modular proteins by small-angle scattering. European Biophysics Journal 39, 769–780 (2010).

39. Kikhney, A. G., Borges, C. R., Molodenskiy, D. S., Jeffries, C. M. & Svergun, D. I. SASBDB: Towards an automatically curated and validated repository for biological scattering data. Protein Science 29, 66–75 (2020).

40. Mastronarde, D. N. Automated electron microscope tomography using robust prediction of specimen movements. J Struct Biol 152, 36–51 (2005).

41. Zivanov, J., Nakane, T. & Scheres, S. H. W. Estimation of high-order aberrations and anisotropic magnification from cryo-EM data sets in RELION-3.1. IUCrJ 7, 253–267 (2020).

42. Rohou, A. & Grigorieff, N. CTFFIND4: Fast and accurate defocus estimation from electron micrographs. J Struct Biol 192, 216–221 (2015).

43. Kimanius, D., Dong, L., Sharov, G., Nakane, T. & Scheres, S. H. W. New tools for automated cryo-EM single-particle analysis in RELION-4.0. Biochemical Journal 478, 4169–4185 (2021).

44. Afonine, P. v et al. Real-space refinement in PHENIX for cryo-EM and crystallography. Acta Crystallographica Section D 74, 531–544 (2018).

45. Emsley, P., Lohkamp, B., Scott, W. G. & Cowtan, K. Features and development of Coot. Acta Crystallogr D Biol Crystallogr 66, 486–501 (2010).

46. Pettersen, E. F. et al. UCSF Chimera—A visualization system for exploratory research and analysis. J Comput Chem 25, 1605–1612 (2004).

47. Pettersen, E. F. et al. UCSF ChimeraX: Structure visualization for researchers, educators, and developers. Protein Science 30, 70–82 (2021).

48. Jumper, J. et al. Highly accurate protein structure prediction with AlphaFold. Nature 596, 583–589 (2021).

49. Hu, G. et al. flDPnn: Accurate intrinsic disorder prediction with putative propensities of disorder functions. Nat Commun 12, (2021).

50. Xue, B., Dunbrack, R. L., Williams, R. W., Dunker, A. K. & Uversky, V. N. PONDR-FIT: A Meta-Predictor of Intrinsically Disordered Amino Acids. Biochim Biophys Acta 1804, 996–1010 (2010).

51. Erdős, G. & Dosztányi, Z. Analyzing Protein Disorder with IUPred2A. Curr Protoc Bioinformatics 70, e99 (2020).

52. Erdős, G., Pajkos, M. & Dosztányi, Z. IUPred3: prediction of protein disorder enhanced with unambiguous experimental annotation and visualization of evolutionary conservation. Nucleic Acids Res 49, W297–W303 (2021).

53. Bernadó, P. Effect of interdomain dynamics on the structure determination of modular proteins by small-angle scattering. European Biophysics Journal 39, 769–780 (2010).

54. Kikhney, A. G. & Svergun, D. I. A practical guide to small angle X-ray scattering (SAXS) of flexible and intrinsically disordered proteins. FEBS Letters vol. 589 2570–2577 Preprint at https://doi.org/10.1016/j.febslet.2015.08.027 (2015).

55. Durand, D. et al. NADPH oxidase activator p67phox behaves in solution as a multidomain protein with semi-flexible linkers. J Struct Biol 169, 45–53 (2010).

56. Bernadó, P., Mylonas, E., Petoukhov, M. v., Blackledge, M. & Svergun, D. I. Structural characterization of flexible proteins using small-angle X-ray scattering. J Am Chem Soc 129, 5656–5664 (2007).

57. Tria, G., Mertens, H. D. T., Kachala, M. & Svergun, D. I. Advanced ensemble modelling of flexible macromolecules using X-ray solution scattering. IUCrJ 2, 207–217 (2015).

58. Ranatunga, W., Jackson, D., Flowers, R. A. & Borgstahl, G. E. O. Human RAD52 protein has extreme thermal stability. Biochemistry 40, 8557–8562 (2001).

59. Nwanochie, E. & Uversky, V. N. Structure determination by single-particle cryo-electron microscopy: Only the sky (and intrinsic disorder) is the limit. International Journal of Molecular Sciences vol. 20 Preprint at https://doi.org/10.3390/ijms20174186 (2019).

60. Greenfield, N. J. Analysis of circular dichroism data. Methods Enzymol 383, 282–317 (2004).

61. Kelly, S. M., Jess, T. J. & Price, N. C. How to study proteins by circular dichroism. Biochimica et Biophysica Acta (BBA) - Proteins and Proteomics 1751, 119–139 (2005).

62. Tompa, P. Intrinsically disordered proteins: a 10-year recap. Trends Biochem Sci 37, 509–516 (2012).

63. Mollica, L. et al. Binding mechanisms of intrinsically disordered proteins: Theory, simulation, and experiment. Frontiers in Molecular Biosciences vol. 3 Preprint at https://doi.org/10.3389/fmolb.2016.00052 (2016).

64. van der Lee, R. et al. Classification of intrinsically disordered regions and proteins. Chemical Reviews vol. 114 6589–6631 Preprint at https://doi.org/10.1021/cr400525m (2014).

65. Park, M. S., Ludwig, D. L., Stigger, E. & Lee, S.-H. Physical Interaction between Human RAD52 and RPA Is Required for Homologous Recombination in Mammalian Cells*.

66. Keith Dunker, A. J. Brown, C., David Lawson, J. M. Iakoucheva, L. & Obradović, Z. Intrinsic Disorder and Protein Function. Biochemistry 41, 6573–6582 (2002).

67. Ma, C. J., Kwon, Y., Sung, P. & Greene, E. C. Human RAD52 interactions with replication protein a and the RAD51 presynaptic complex. Journal of Biological Chemistry 292, 11702–11713 (2017).

