## Supplemental Figures 1-11; Tables 1-3 for "Novel structural insights on full-length human RAD52: Cryo-EM and beyond"

**Supplementary Figures**

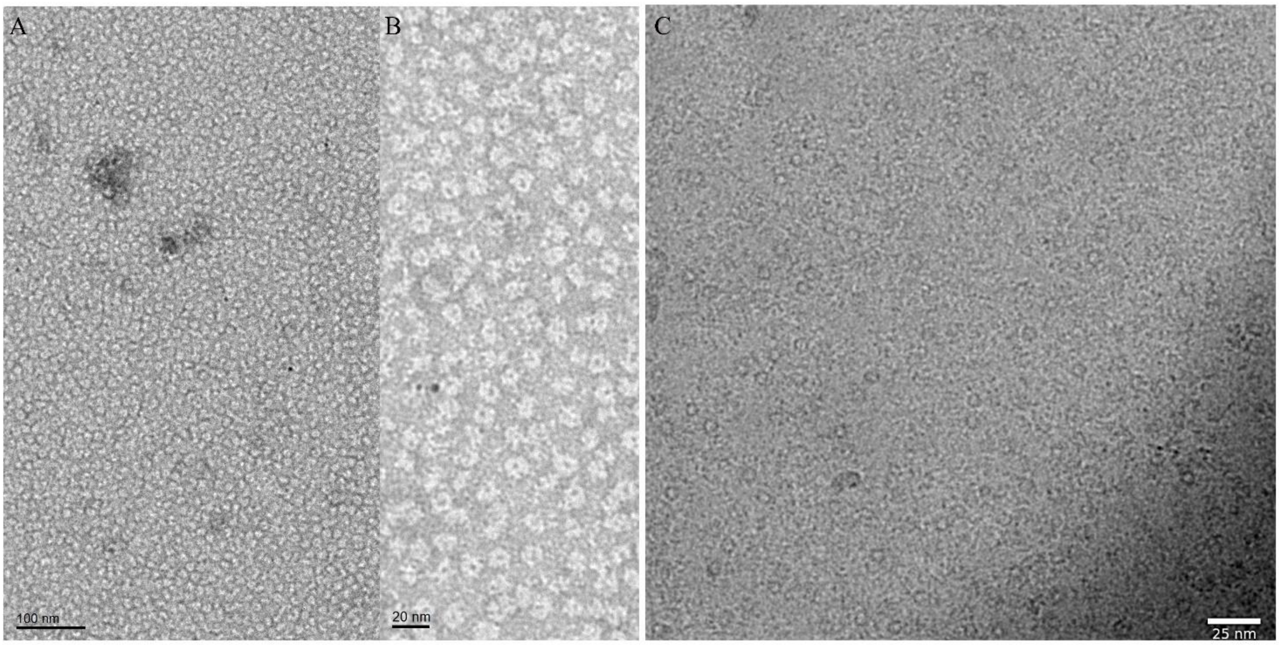


Supplementary Fig. 1) Electron microscopy micrographs of RAD52 FL. A, B: negative stained and C: cryo-EM representative micrographs.

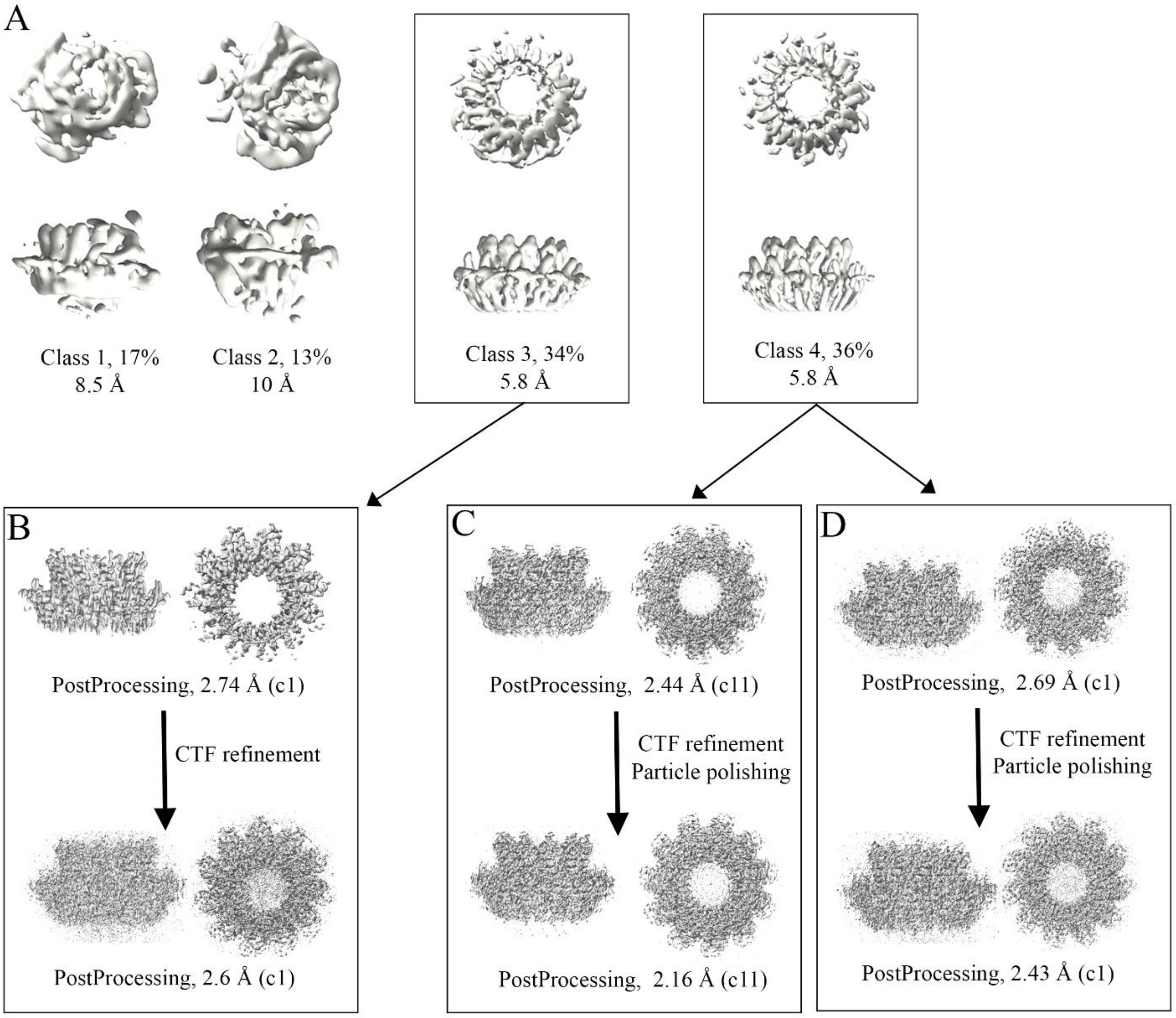


Supplementary Fig. 2) RAD52 FL high-resolution analysis. A: RAD52 FL 3D class averages in top and side views based on 2325722 particles with their rlnClassDistribution and rlnResolution; B: RAD52 FL cryo-EM density maps obtained from 3D class 3 imposing c1 symmetry after CTF refinement; C: RAD52 FL cryo-EM density maps obtained from 3D class 4 imposing c11 symmetry after CTF refinement and particle polishing; D: RAD52 FL cryo-EM density maps obtained from 3D class 4 imposing no symmetry (c1) after CTF refinement and particle polishing.

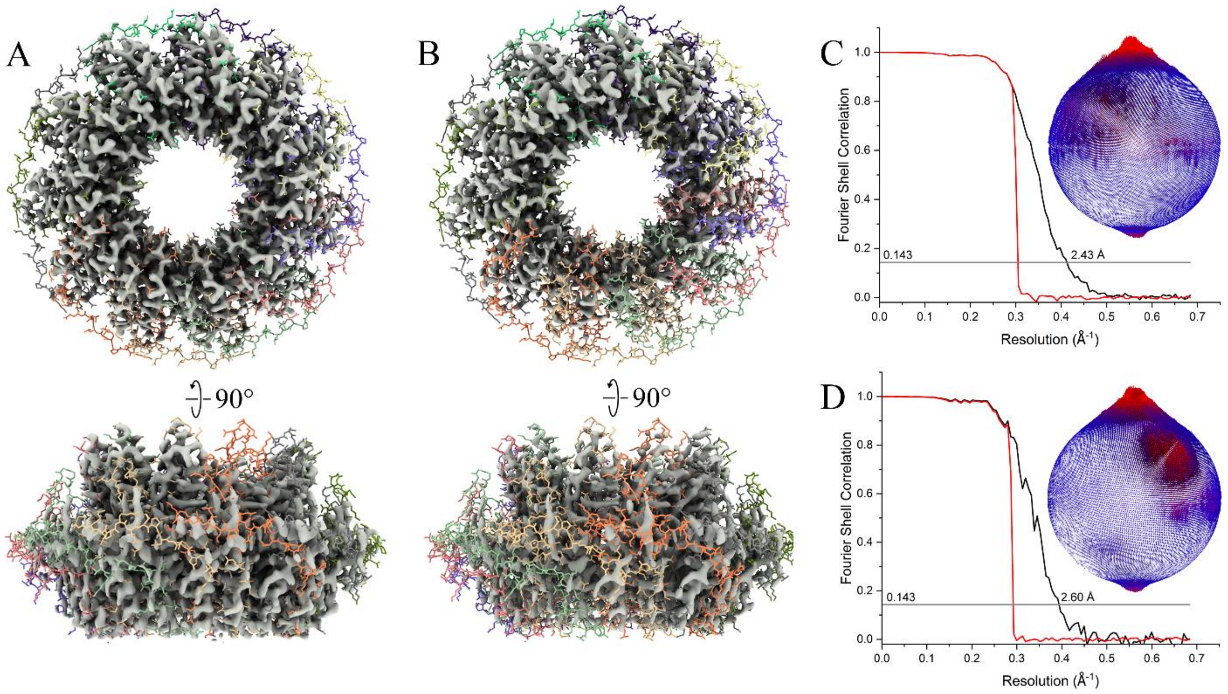


Supplementary Fig. 3) A: RAD52 FL cryo-EM density maps obtained from 3D class 4 without imposing symmetry (c1) after CTF refinement and particle polishing; B: RAD52 FL cryo-EM density maps obtained from 3D class 3 without imposing symmetry (c1) after CTF refinement. C: FSC curve of the RAD52 FL of cryo electron density map in A; D: FSC curve of the RAD52 FL of cryo electron density map in B.

A

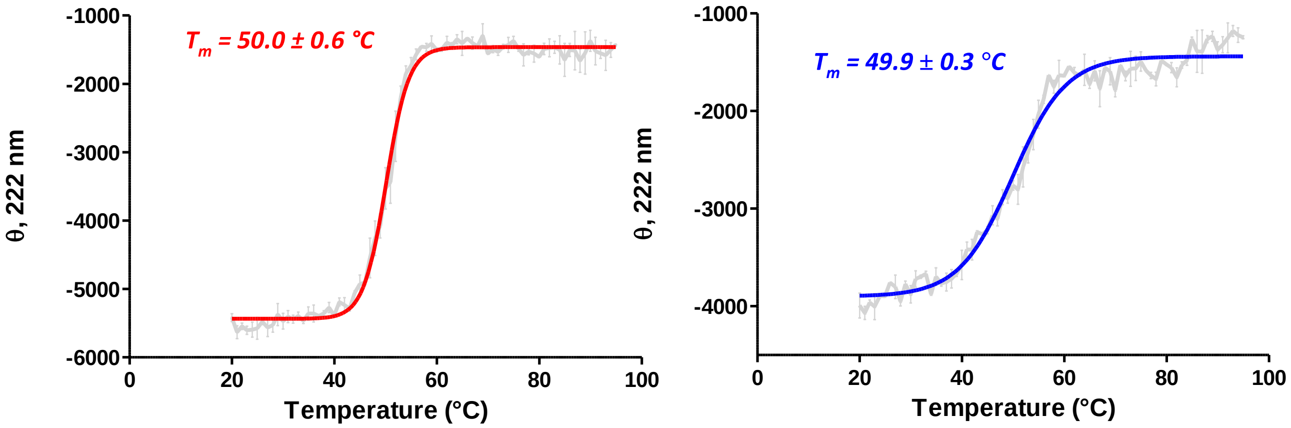


B

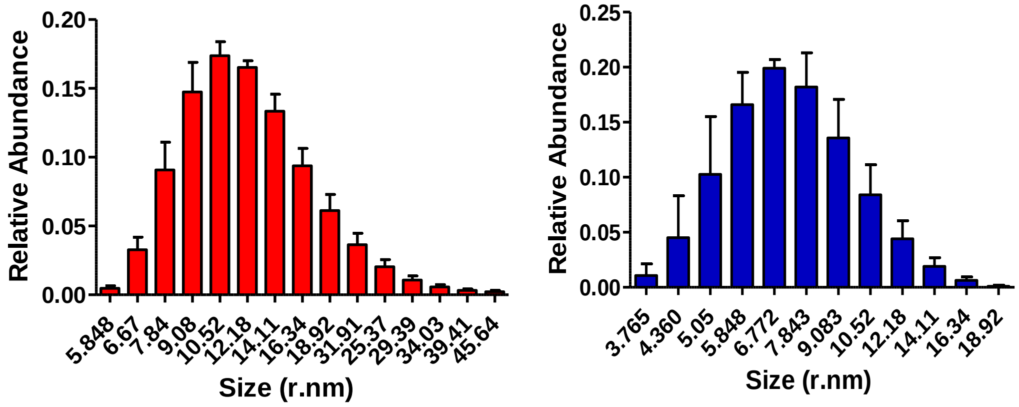


Supplementary Fig. 4) RAD52 FL and N-terminal domain stability and superstructures analyses A: Thermal stability analysis: comparison of θ at 222 nm of RAD52 FL (red) and RAD52 N-terminal (blue) for melting temperature ($T_m$) determination; data analyses were performed after normalization; B: DLS profile of RAD52 FL protein size analysis (red histogram) shows a heterogeneous protein sample with high molecular weight RAD52 superstructures, which are on average larger compared to the ones observed in the DLS profile of RAD52 N-terminal domain protein sample (blue histogram).

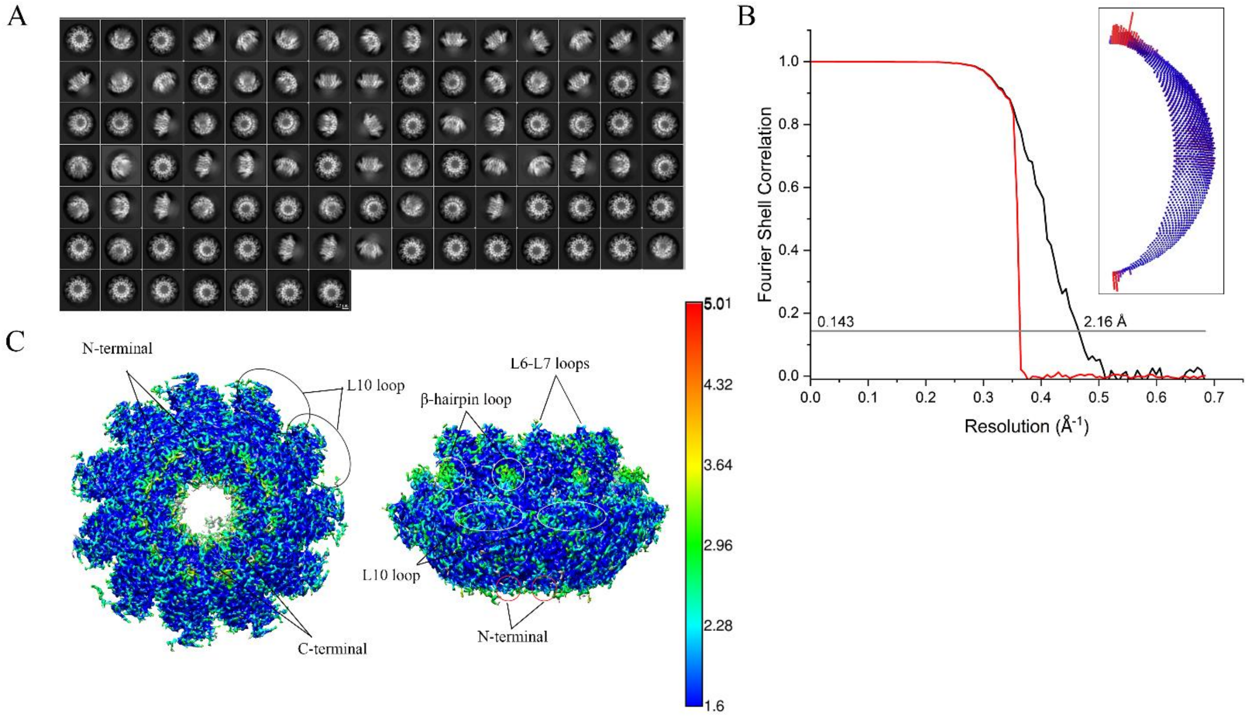


Supplementary Fig. 5) A: representative unsupervised RAD52 FL 2D class averages of 1616139 particles. B: FSC curve of the RAD52 FL (red, FSC phase randomized masked curve; black, FSC corrected curve) with a resolution corresponding to FSC=0.143 marked. The inset shows the Euler angle distribution. C: RAD52 FL cryo-EM electron density map filtered according to ResMap local resolution.

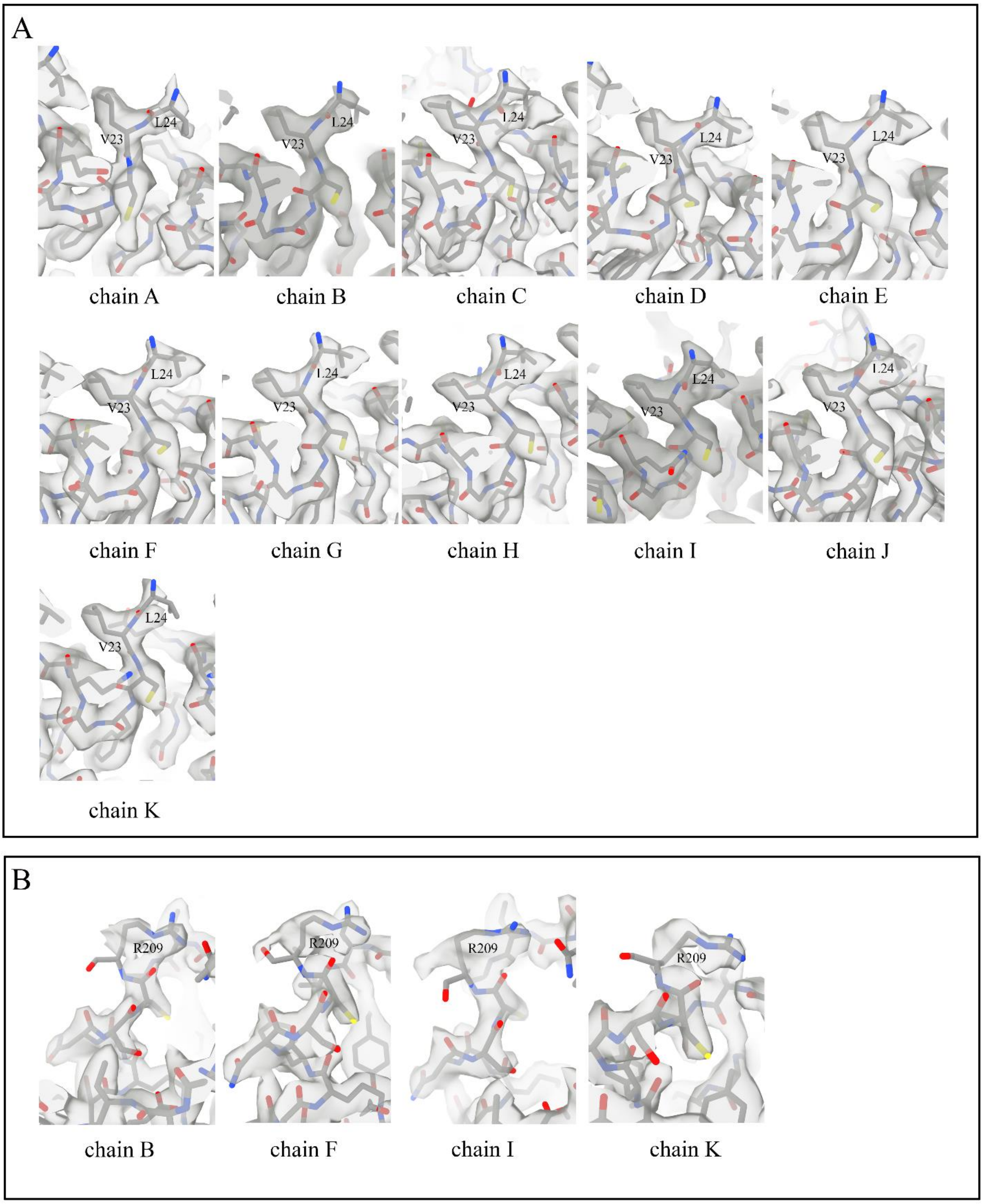


Supplementary Fig. 6) Detail of Val 23 and Leu24 residues at the N-terminal (A) and Arg209 residues at the C-terminal of the RAD52 FL model. The model is fitted in its corresponding cryo-EM electron density map.

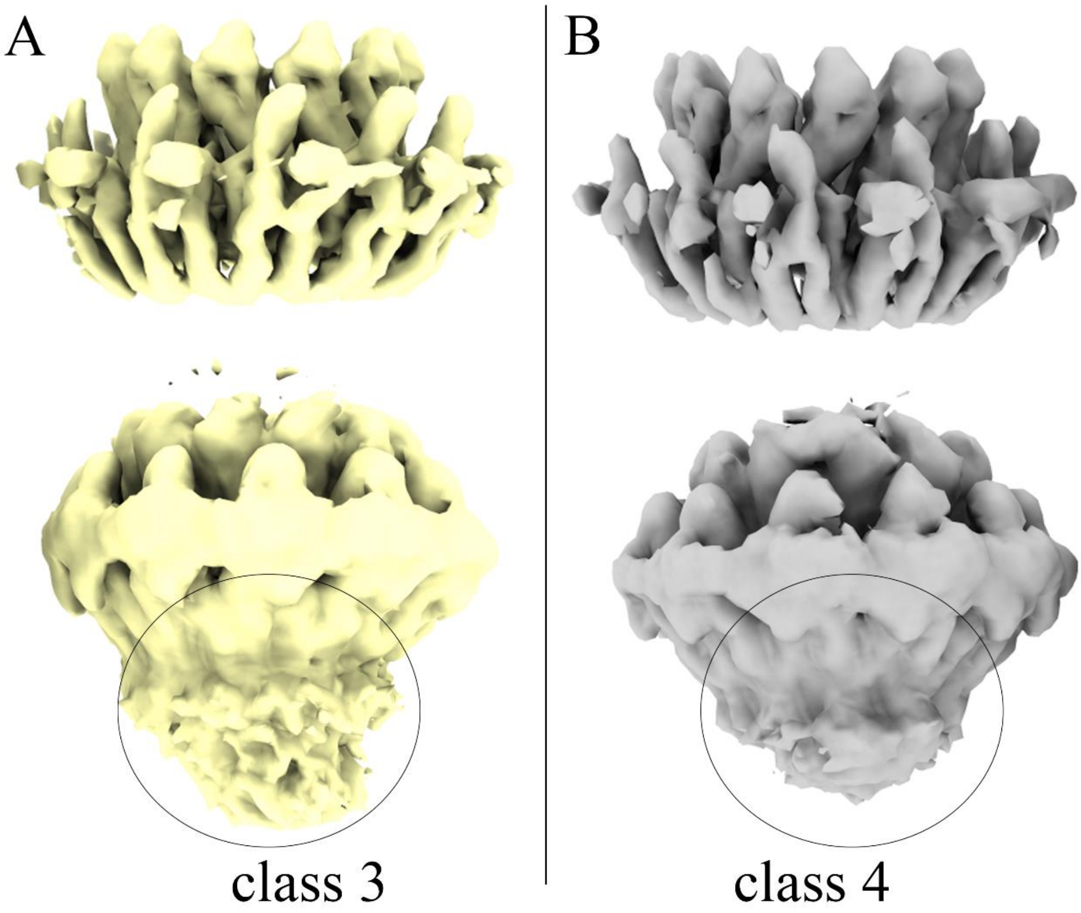


Supplementary Fig. 7) A: Cryo-EM electron density map of RAD52 FL 3D class averages number 3 in side views shown at a higher (up) and lower (bottom) density threshold; B: Cryo-EM electron density map of RAD52 FL 3D class averages number 4 in side views shown at a higher (up) and lower (bottom) density threshold. Note the presence of an unstructured region close to the top of the ring (encircled).

A

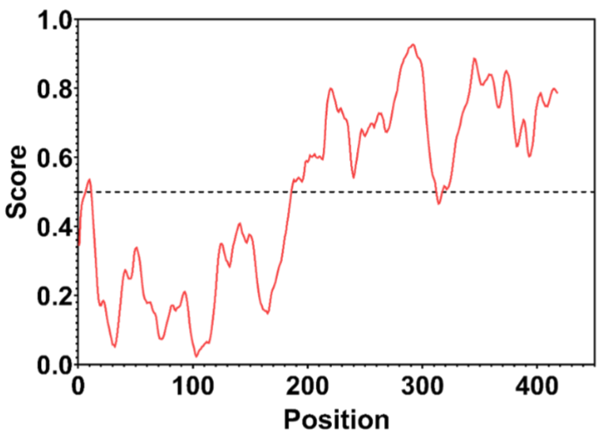
Score
Position
0.0
0.2
0.4
0.6
0.8
1.0
0
100
200
300
400

B

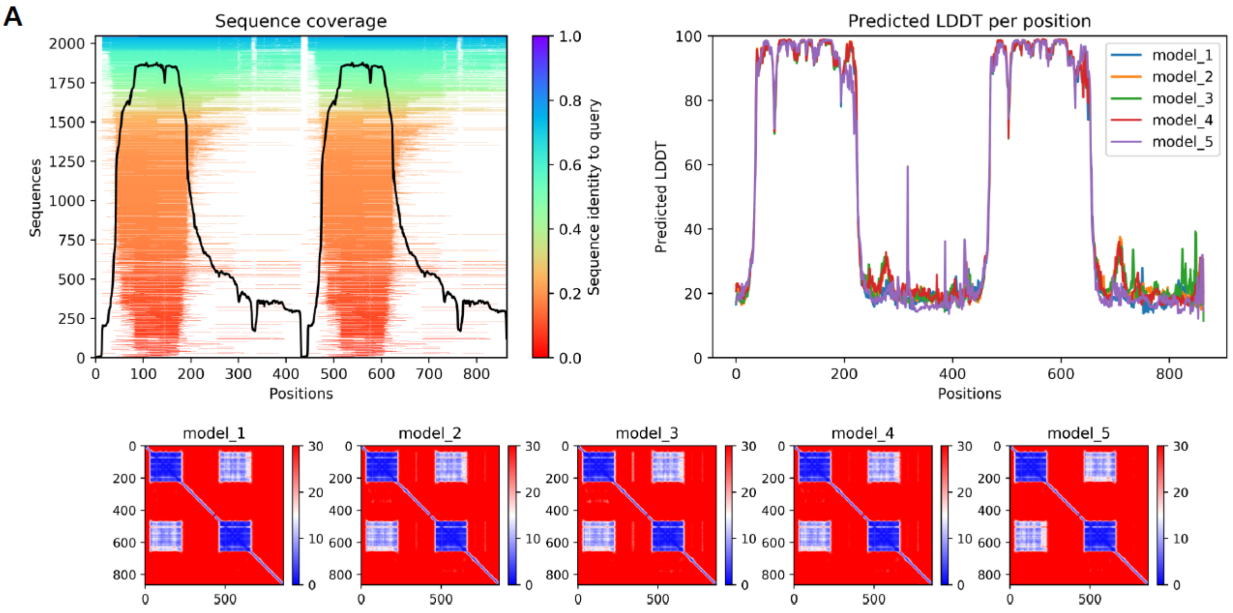
A
Sequence coverage
Sequences
Positions
Sequence identity to query
Predicted LDDT per position
Predicted LDDT
model_1
model_2
model_3
model_4
model_5

C

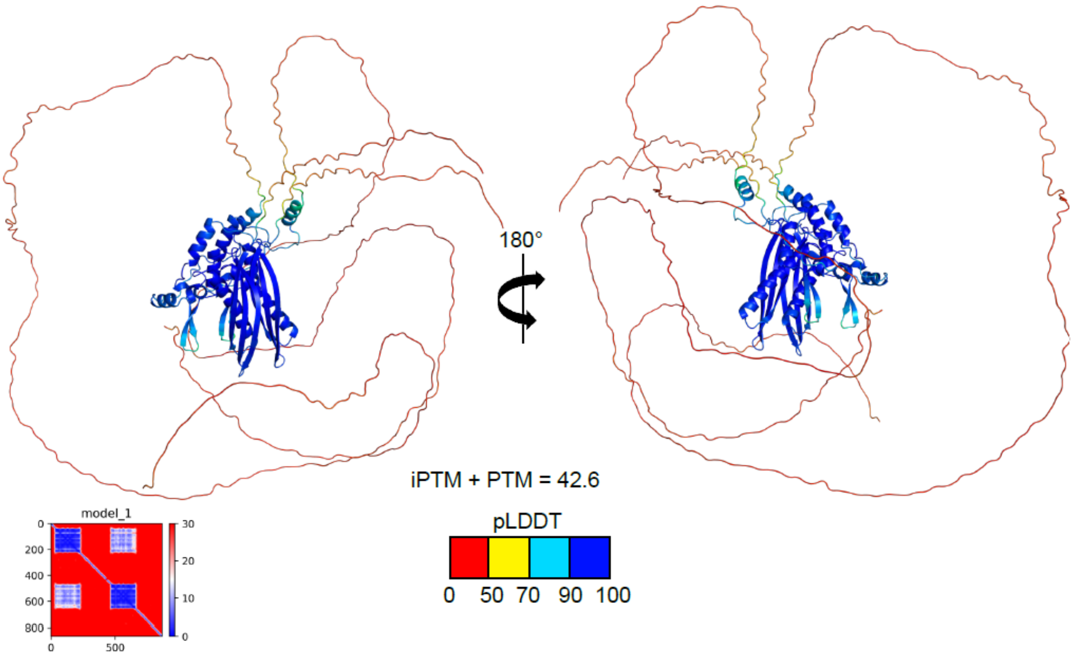
180°
iPTM + PTM = 42.6
model_1
pLDDT
0
50
70
90
100

Supplementary Fig. 8) A: IUPRED analyses on RAD52 FL sequence identifying the C-terminal domain of RAD52 as an intrinsically disordered region (IUPRED score >0.5), B: Left: Multi Sequence Analysis (MSA) depth and diversity. Right: AlphaFold2 confidence predicted local distance difference test (pLDDT). Bottom: Predicted Align Error; C: structure of the Alphafold model ranked with the highest average Score, interface predicted template modelling score + predicted modelling score (iPTM + PTM). Residues have been colored for local model confidence. Bottom left: predicted aligned error (PAE) of the corresponding model.

A

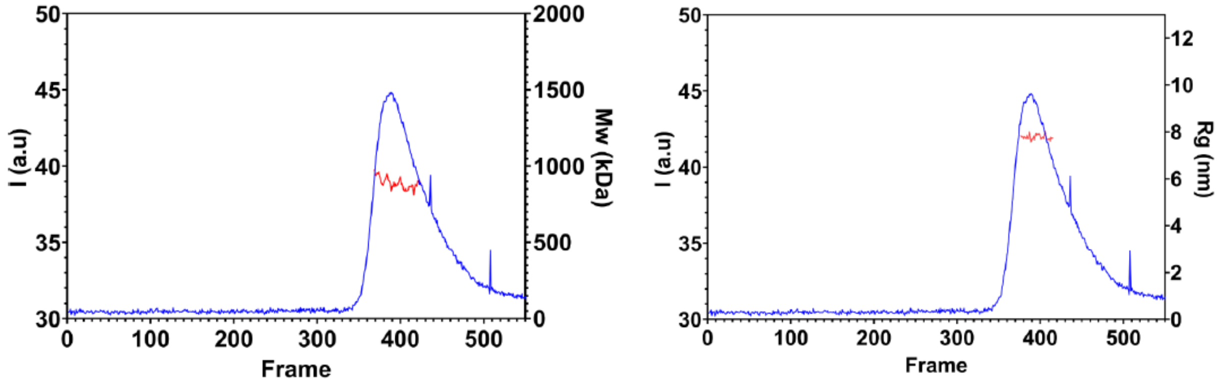


B

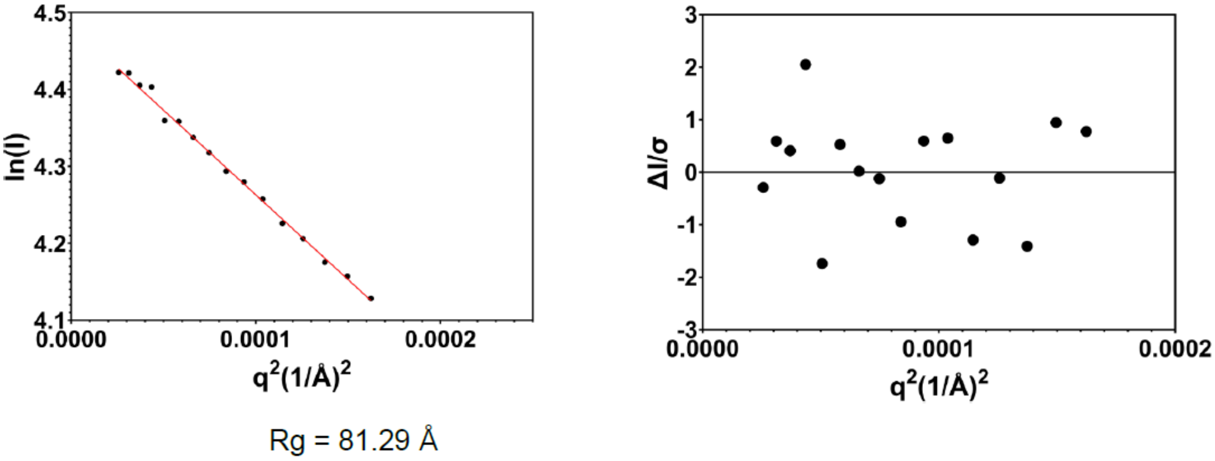


Supplementary Fig. 9) A: SEC - SAXS profile of RAD52 FL in 20 mM Tris pH 7.5, 250 mM NaCl, 1% glycerol buffer, left average of mw, right average of Radius of Gyration; B: Guiner analysis of SAXS Profile. On the left Guiner fitting, on the right residuals of Guiner fitting.

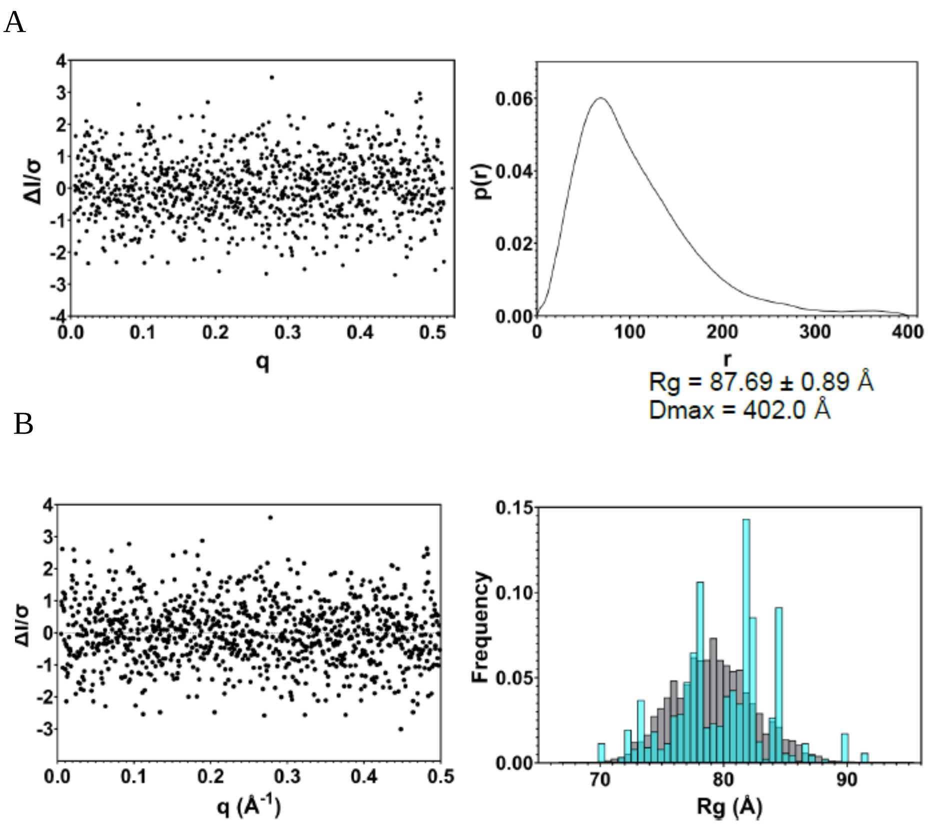


Supplementary Fig. 10) Small angle X-ray scattering analysis of RAD52 FL (1.3 mg/mL); A: SAXS p(r) analysis. Left: fitting residuals; Right: p(r) plot and calculated Dmax; B: EOM Modelling of the N-terminal and C-terminal domain of RAD52FL utilizing as a rigid body the Cryo-EM structure. Left: model residuals. Right: comparison of distribution of radius of gyration in selected models (cyan) and generated pool (grey).

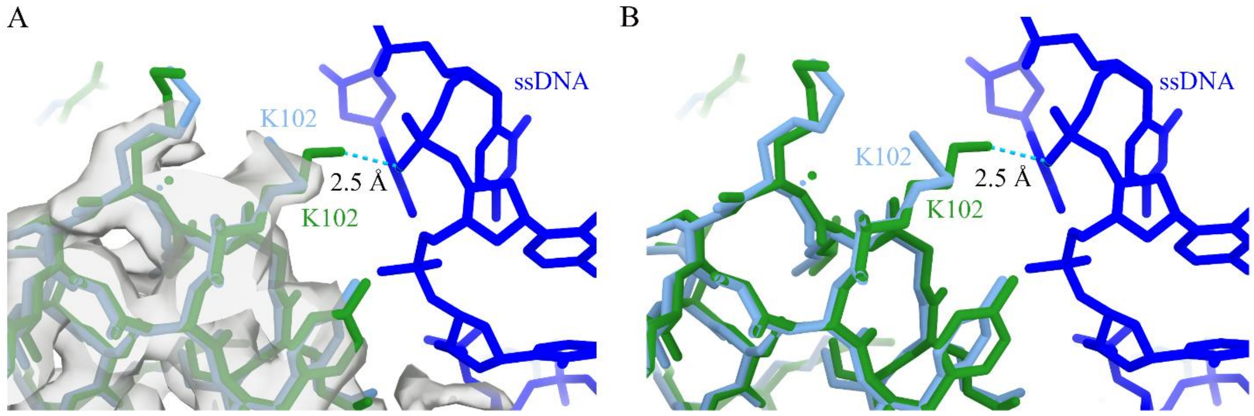


Supplementary Fig. 11) Detail of the RAD52 FL model region containing Lys102 (in cyan) and fitted in the cryo-EM electron density map. The model is superimposed to the same region from the crystal structure of the $RAD52_{25-208}$ outer DNA binding site model (PDB ID: 5XS0[28]) in green. Note that the cryo-electron density map is shown at lower density threshold.

Supplementary Table 1

(a) Sample details

| | |
|---|---|
| Organism | Homo sapiens (Human) |
| Source | *E. Coli* expressed |
| Uniprot sequence ID | P43351 |
| Extinction coefficient [$A_{280}$, 0.1%(w/v)] | 0.873 |
| $\bar{v}$ from chemical composition of N-terminal domain (PDB entry : 1KN0) ($cm^3$/ g) | 0.73 $cm^3$/g |
| Particle contrast from sequence and solvent constituents $\bar{\rho}$ ($\rho_{protein}$ - $\rho_{solvent}$) $10^{-6}$ $Å^{-2}$ | 2.503 |
| M from chemical composition (Da) | 527540 |
| SEC–SAXS column | Superose™ 6 Increase 3.2/300 |
| Loading concentration (mg/mL) | 1.3 |
| Injection Volume (mL) | 100 µL |
| Flow rate (mL/min) | 0.075 mL/min |
| Solvent (solvent blanks taken from SEC flow-through prior to elution of protein) | 25 mM Tris pH 7.5, 250 mM NaCl, 1% Glycerol |

(b) SAXS data-collection parameters

| | |
|---|---|
| Instrument/data processing | FreeSAS[1] |
| Wavelength (Å) | 0.99 (12.5KeV) |
| Beam size at sample (mm) | 0.200 × 0.100 at sample plane |
| Beam size at detector (focus) (mm) | 1 pixel ~ 0.1 x 0.1 |
| Camera length (m) | 2.81 |
| q measurement range (Å) | 0.007–0.55 |
| Absolute scaling method | Water |
| Sample configuration | SEC-SAXS |
| Sample temperature (°C) | 20 |
| Exposure time | 2 second / frame (600 frames) |

(c) Software employed for SAXS data reduction, analysis and interpretation

| | |
|---|---|
| SAXS data reduction | FreeSAS[1] Solvent subtraction and frame selection were performed using Chromixs (ATSAS 3.1.1) [2,3] |
| Extinction coefficient estimate | Expasy ProtParam tool [4] |
| Calculation of $\bar{v}$ and $\bar{\rho}$ values | Biomolecular Scattering Length Density Calculator (http://psldc.isis.rl.ac.uk/Psldc/ ) |
| Basic analyses: Guinier, P(r), VP | BioXTAS RAW, GNOM [5,6] |
| Atomic structure modelling | RANCH via ATSAS Online (https://www.embl-hamburg.de/biosaxs/atsas-online/)[7,8]<br>FFMAKER, ATSAS 3.1.1 (run on local Pc)[7,8]<br>GAJOE 2.1, ATSAS 3.1.1 (run on local Pc)[7,8] |

(d) Structural parameters

| | |
|---|---|
| Guinier analysis | |
| I(0) ($cm^{-1}$) | 88.51 ± 0.34 |
| $R_g$ (Å) | 81.29 |
| $q_{min}$ ($Å^{-1}$) | 0.00507 |
| $qR_g$ max | 1.04 |
| Coefficient of correlation, $R^2$ | 0.9957 |
| P(r) analysis | |
| I(0) ($cm^{-1}$) | 89.86 ± 0.36 |
| $R_g$ (Å) | 87.69 ± 0.89 |
| dmax (Å) | 402.0 |
| q range ($Å^{-1}$) | 0.0051 – 0.5156 |
| $\chi^2$ (total estimate from GNOM) | 0.9343 |
| GNOM Interpretation | A Reasonable Solution |
| Porod volume ($Å^{-3}$) | 1330000 |
| V, M using the Fischer method | 818000, 678.7 kDa |
| $V_c$, M | 592.9 kDa |
| Bayesian inference | 585.2 kDa |

(e) Atomistic modelling.

| Multistate/ensemble models | |
|---|---|
| Ensemble Optimization Method [7,8] (default parameters, 20000 models in initial ensemble, disordered models, constant subtraction, curve repetition in ensemble, maximum number of curves per ensemble 20, minimum number of curves per ensemble 5, curve repetition in the ensemble allowed, number of cycles of the genetic algorithm to run (min. 1): 100 | |
| Structures | PDB Entry : 8BJM |
| q range for all modelling (Å) | 0.0507 – 0.5 |
| $\chi^2$, CORMAP P-value | 0.960, 0.208 |
| Constant subtraction | 0.058 |
| No. of representative structures | 4 |

(f) SASBDB IDs for data and models[9]

| | |
|---|---|
| His-RAD52 FL | SASDQ49 – His-Tagged full length DNA repair protein RAD52 homolog |

(g) ESRF - DOI

All data are saved in hierarchical data format (HDF) will be available at [10]


1. Tully, M. D. *et al.* BioSAXS at European Synchrotron Radiation Facility – Extremely Brilliant Source: BM29 with an upgraded source, detector, robot, sample environment, data collection and analysis software. *J. Synchrotron Radiat.* **30**, 1–9 (2023).

2. Panjkovich, A. & Svergun, D. I. CHROMIXS: automatic and interactive analysis of chromatography-coupled small-angle X-ray scattering data. *Bioinformatics* **34**, 1944–1946 (2018).

3. Manalastas-Cantos, K. *et al.* ATSAS 3.0 : expanded functionality and new tools for small-angle scattering data analysis. *J. Appl. Crystallogr.* **54**, 343–355 (2021).

4. Gasteiger, E. *et al.* The Proteomics Protocols Handbook. *Proteomics Protoc. Handb.* 571–608 (2005)

doi:10.1385/1592598900.

5. Hopkins, J. B., Gillilan, R. E. & Skou, S. BioXTAS RAW: Improvements to a free open-source program for small-angle X-ray scattering data reduction and analysis. *J. Appl. Crystallogr.* **50**, 1545–1553 (2017).

6. Svergun, D. I. Determination of the regularization parameter in indirect-transform methods using perceptual criteria. *J. Appl. Crystallogr.* **25**, 495–503 (1992).

7. Tria, G., Mertens, H. D. T., Kachala, M. & Svergun, D. I. Advanced ensemble modelling of flexible macromolecules using X-ray solution scattering. *IUCrJ* **2**, 207–217 (2015).

8. Bernadó, P., Mylonas, E., Petoukhov, M. V., Blackledge, M. & Svergun, D. I. Structural characterization of flexible proteins using small-angle X-ray scattering. *J. Am. Chem. Soc.* **129**, 5656–5664 (2007).

9. Kikhney, A. G., Borges, C. R., Molodenskiy, D. S., Jeffries, C. M. & Svergun, D. I. SASBDB: Towards an automatically curated and validated repository for biological scattering data. *Protein Sci.* **29**, 66–75 (2020).

10. Rinaldi, F., Hočevar, J. & Scietti, L. Proteins related to pathogenesis of diseases, viral proteins, cell division, signalling and chromatin processes [Dataset]. *European Synchrotron Radiation Facility* (2025) doi:https://doi.org/10.15151/ESRF-ES-771426690.

Supplementary Table 2

DATA COLLECTION

| | Screening session | High-resolution Session |
|---|---|---|
| Microscope model | Thermo Fisher Scientific Glacios Selectris X | Thermo Fisher Scientific IC-Krios Selectris X |
| Detector type | Thermo Fisher Scientific Falcon 4 EC | Thermo Fisher Scientific Falcon 4 EC |
| Imaging mode | EF-TEM | EF-TEM |
| Accelerating voltage, kV | 200 | 300 |
| Pixel size, Å | 1.154 Å/pix | 0.731 Å/pix |
| Total exposure time, sec | 8.6 sec | 8 $e^-$/pix/sec |
| Total Number of collected stacks | 831 | 17400 |
| Number of stacks used in the analysis | 831 | 17400 |
| Total dose per stack, e−/Å2 | 40.9 $e^-/Å^2$ | 50 $e^-/Å^2$ |
| Number of frames per stack | 34 | 49 |
| Defocus range, µm | -1.2 um to -2.5 um | -0.8 um to -1.8 |

DATA PROCESSING, GLOBAL RESOLUTION (Å), PDB ID AND EMDB ID

| | Screening session | High-resolution Session |
|---|---|---|
| 3D reconstruction software package | Relion 3.1 | Relion 4.0 |
| Extracted particles | 580952 | 2325722 |
| Refined particles | 221976 | 837272 |
| Symmetry | C11 | C11 |
| FSC0.143 (unmasked/masked) | 3.4 Å/pix | 2.16 Å/pix |
| PDB ID | - | 8BJM |
| EMBD ID | - | EMD-16089 |

Supplementary Table 3

| **hRAD52 Models** | **RMSD** |
|---|---|
| hRAD52$_{25-208}$ (PDB ID: 1KN0) | 0.625 |
| hRAD52$_{25-208}$ inner DNA binding site (PDB ID: 5XRZ) | 0.547 |
| hRAD52$_{25-208}$ outer DNA binding site (PDB ID: 5XS0) | 0.539 |

Root-mean-square deviation (RMSD) between the hRAD52 FL cryoEM model and the RAD52$_{25-208}$ crystallographic models
